# The variation and evolution of complete human centromeres

**DOI:** 10.1101/2023.05.30.542849

**Authors:** Glennis A. Logsdon, Allison N. Rozanski, Fedor Ryabov, Tamara Potapova, Valery A. Shepelev, Yafei Mao, Mikko Rautiainen, Sergey Koren, Sergey Nurk, David Porubsky, Julian K. Lucas, Kendra Hoekzema, Katherine M. Munson, Jennifer L. Gerton, Adam M. Phillippy, Ivan A. Alexandrov, Evan E. Eichler

**Author notes:** **Correspondence to:** Evan E. Eichler, Ph.D. Department of Genome Sciences, University of Washington School of Medicine 3720 15th Ave NE, S413A, Seattle, WA 98195-5065. Institute for Molecular Medicine Finland (FIMM), Helsinki Institute of Life Science (HiLIFE), University of Helsinki, Helsinki, Finland.

## Abstract

We completely sequenced and assembled all centromeres from a second human genome and used two reference sets to benchmark genetic, epigenetic, and evolutionary variation within centromeres from a diversity panel of humans and apes. We find that centromere single-nucleotide variation can increase by up to 4.1-fold relative to other genomic regions, with the caveat that up to 45.8% of centromeric sequence, on average, cannot be reliably aligned with current methods due to the emergence of new α-satellite higher-order repeat (HOR) structures and two to threefold differences in the length of the centromeres. The extent to which this occurs differs depending on the chromosome and haplotype. Comparing the two sets of complete human centromeres, we find that eight harbor distinctly different α-satellite HOR array structures and four contain novel α-satellite HOR variants in high abundance. DNA methylation and CENP-A chromatin immunoprecipitation experiments show that 26% of the centromeres differ in their kinetochore position by at least 500 kbp—a property not readily associated with novel α-satellite HORs. To understand evolutionary change, we selected six chromosomes and sequenced and assembled 31 orthologous centromeres from the common chimpanzee, orangutan, and macaque genomes. Comparative analyses reveal nearly complete turnover of α-satellite HORs, but with idiosyncratic changes in structure characteristic to each species. Phylogenetic reconstruction of human haplotypes supports limited to no recombination between the p- and q-arms of human chromosomes and reveals that novel α-satellite HORs share a monophyletic origin, providing a strategy to estimate the rate of saltatory amplification and mutation of human centromeric DNA.

## INTRODUCTION

Advances in long-read sequencing technologies and assembly algorithms have now enabled the complete assembly of complex repetitive regions in the human genome, including centromeres, for the first time^1–5^. In addition to these technological advances, completion of the first human genome was aided by the use of a complete hydatidiform mole (CHM)^3^—an abnormality of development where only the paternal chromosomal complement is retained. The particular cell line, CHM13, simplified the assembly process because the presence of a single human haplotype eliminated allelic variation that can otherwise complicate the assembly of structurally complex regions^1, 6^. This combination of technologies and resources, thus, provided the first complete sequence of each centromere from a single human genome^3, 4^. Notwithstanding these advances, human centromeres still pose a challenge to sequencing and assembly. In a recent analysis of human genomes sequenced as part of the Human Pangenome Reference Consortium (HPRC), no other human genome was completely sequenced across its centromeres^7^. The centromeres, in particular, were among the most gap-ridden regions^8^ and excluded from the construction of a pangenome^7^. Additional methods and approaches are still required to fully sequence and assemble these regions^9^.

Human centromeres have been shown to represent some of the most diverse and rapidly evolving regions in the genome. The bulk of human centromeric DNA is composed of tandemly repeating, ∼171-bp α-satellite DNA, which are organized into higher-order repeat (HOR) units that can extend for megabase pairs (Mbp) of sequence and are particularly variable among humans due to the action of unequal crossing over, concerted evolution, and saltatory amplification. Thus, a single human genome, such as CHM13, cannot adequately represent or capture human genetic diversity. While most of the human genome has been interrogated for allelic variation at the base-pair level, studies of centromeric DNA are far more limited, based on early pulsed-field gels and Southern blots^10–12^, monomer α-satellite analyses of short reads^13, 14^, or analyses restricted to select regions or chromosomes^4, 15, 16^. Here, we set out to sequence a complete set of centromeres from another human genome using a second hydatidiform mole cell line (CHM1)^1, 6, 17^. We compare two complete sets of human centromeres to establish a baseline for single-nucleotide and structural variation. We relate these differences to shifts in the sites of kinetochore attachment and compare the rate and tempo of mutational change of centromeric DNA by sequencing select chromosomes from other nonhuman primate (NHP) species and comparing our findings to finished centromeres from the HPRC^7^ and Human Genome Structural Variation Consortium (HGSVC)^18^.

## RESULTS

### Complete sequence and validation of CHM1 centromeres

To assemble each centromere in the CHM1 genome, we developed an approach similar to that used for the assembly of the CHM13 centromeres (**Extended Data Fig. 1**). First, we generated ∼66-fold sequence coverage of Pacific Biosciences (PacBio) high-fidelity (HiFi) sequence data and ∼98-fold coverage of Oxford Nanopore Technologies (ONT) data from the complete hydatidiform mole cell line CHM1 (**Extended Data Table 1**). We initially used the whole-genome assembler hifiasm^19^ to generate a highly accurate backbone genome assembly. Only four centromeres were contiguously assembled (from chromosomes 2, 7, 19, and 20), with the remaining 19 fragmented into multiple contigs. We resolved the remaining centromeres by using singly unique nucleotide *k*-mers (SUNKs) to barcode the PacBio HiFi contigs, bridging them with ultra-long (>100 kbp) ONT reads that share a similar barcode, as described previously^16^. Finally, we improved the base accuracy of the assemblies by replacing the ONT sequences with locally assembled PacBio HiFi contigs, which generated complete sequence assemblies of all CHM1 centromeres with an estimated base accuracy >99.9999% (QV>60; **Methods**).

The use of a complete hydatidiform mole cell line poses particular challenges as it can be subject to somatic rearrangement that arise during culture. This is especially a concern for centromeric satellites, which have been regarded by some as among the most mutable and challenging regions of the genome to assemble^20, 21^. We carefully assessed the CHM1 cell line for chromosomal rearrangements (**Extended Data Figs. 2,3, Supplementary Notes 1,2**) and validated the integrity and biological significance of each CHM1 centromere with a series of experiments. First, we mapped native long-read sequencing data generated from the CHM1 genome to each centromere assembly and confirmed the integrity of all chromosomes with two exceptions (**Extended Data Fig. 4, Supplementary Note 2**). Next, we applied an algorithm, VerityMap^22^, that identifies discordant *k*-mers between the centromere assemblies and PacBio HiFi reads and found no evidence of discordance (**Methods**). Third, we applied a method, GAVISUNK^23^, that compares SUNKs in the centromere assemblies to those in the ONT reads generated from the same sample and observed support for each SUNK with orthogonal ONT data (**Extended Data Fig. 5**). Fourth, we compared the sequence of each CHM1 centromere assembly to those generated by an independent and recently released assembler, Verkko^24^, and found that they were highly concordant, with >99.99% sequence identity between each pair (**Extended Data Fig. 6**). Finally, we compared both the CHM1 and CHM13 genomes directly to 56 genomes (112 haplotypes) sequenced as part of the HPRC^7^ and HGSVC^18^. While many of these additional human genomes are not yet completely assembled across the centromeres, 20.9% of human haplotypes match ≥99% to the newly assembled centromeric regions (**Extended Data Table 2, Extended Data Fig. 7**). In fact, we find that 46.9% of these haplotypes are a better match to CHM1 than to CHM13 (**Extended Data Table 2, Extended Data Fig. 7**), confirming the biological relevance of the CHM1 centromeres.

### Genetic variation among human centromeres

The complete assembly of each CHM1 centromere enables, in principle, a comprehensive comparison of centromeric allelic sequence and structure between two human genomes (Fig. 1). In light of the considerable variation between centromeres and the challenge in creating optimal alignments (especially among α-satellite HORs), we analyzed the blocks of monomeric α-satellite DNA in the pericentromere separately from the α-satellite HOR arrays, and we considered three different alignment strategies, including one designed to specifically handle variation in tandem repeats^25^ (**Methods**). We initially compared the centromeres from the CHM1 and CHM13 genomes and then extended our analysis to both complete and incompletely sequenced centromeres from 56 human genomes (**Extended Data Tables 3-6**). Comparison of the CHM1 and CHM13 centromeres revealed that 63.0-71.5% of α-satellite HORs (depending on the chromosome) could be reliably aligned between the two haplotypes (i.e., >90% identity; **Extended Data Table 3**). Extending this analysis to those from 56 diverse human genomes from the HPRC and HGSVC, we found that this drops to 53.2-55.3% (**Extended Data Table 6**), underscoring the considerable variation in these genomes and the emergence of new HOR structures in some human haplotypes but not others. For the portions that could be aligned, the results were comparable among the three methods (**Extended Data Table 3**), and we report the “full contig alignment” statistics with respect to single-nucleotide variation below (**Methods**).

**Figure 1.**
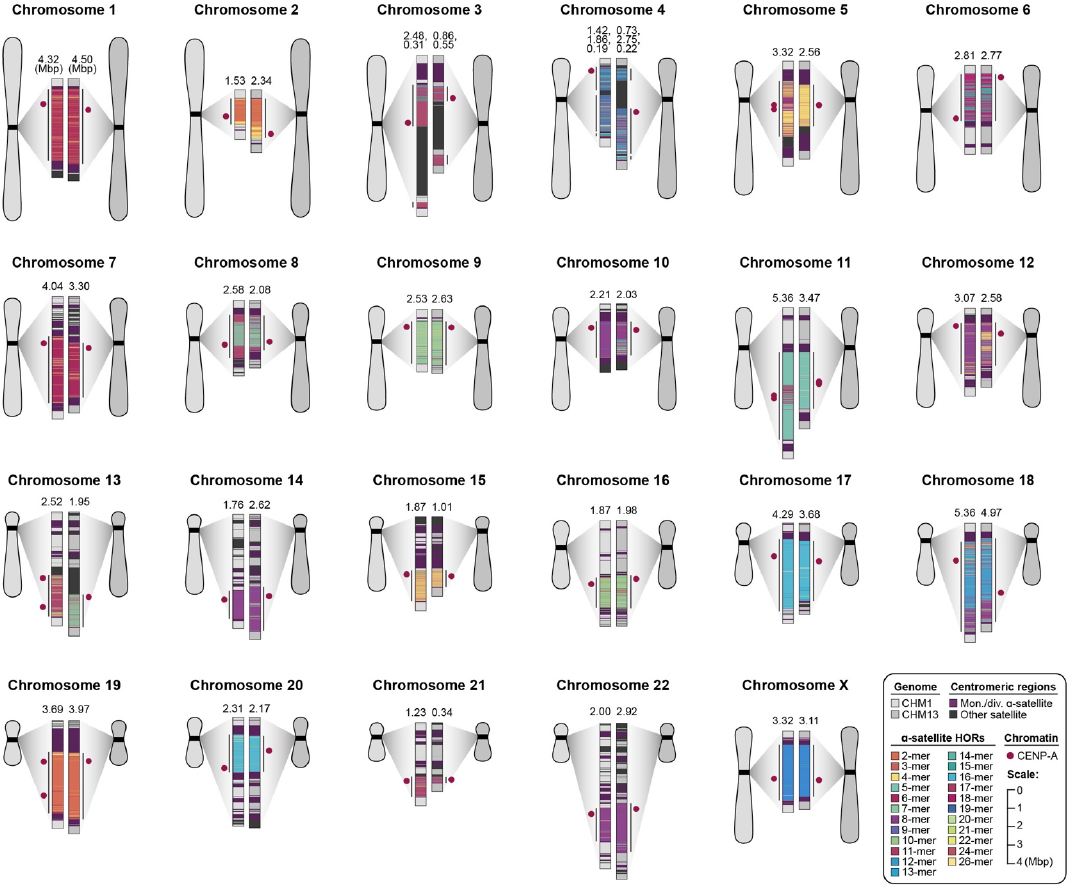
Overview of the centromeric genetic and epigenetic variation between two human genomes. Complete assembly of centromeres from two hydatidiform moles, CHM1 and CHM13, reveals both small- and large-scale variation in centromere sequence, structure, and epigenetic landscape. The CHM1 and CHM13 centromeres are shown on the left and right between each pair of chromosomes, respectively. The length of the α-satellite higher-order repeat (HOR) array(s) is indicated, and the location of centromeric chromatin, marked by the presence of the histone H3 variant CENP-A, is indicated by a dark red circle.

In comparing the CHM1 and CHM13 centromeres to each other, we find that sequence identity increases as we transition from heterochromatin to euchromatin. For example, the mean sequence identity for the alignable portions of CHM1 and CHM13 α-satellite HOR arrays is 98.6 ± 1.6%, in contrast to monomeric/diverged α-satellites at 99.8 ± 0.4% and other pericentromeric satellite DNA (β-satellite, γ-satellite, and human satellites) at 99.1 ± 1.5% (**Extended Data Table 4, Extended Data Fig. 8**). Extending further into the non-satellite pericentromeric DNA, the sequence identity begins to approximate rates of allelic variation (99.9 ± 0.3%; **Extended Data Table 4, Extended Data Fig. 8**). We note, however, that this varies considerably depending on the chromosome (Fig. 2a, **Extended Data Fig. 9**), and the presence of imperfectly aligned α-satellite repeats further complicates such calculations. The centromeres of some chromosomes, such as 19 and X, show the highest degree of concordance between their α-satellite HOR arrays, whereas all others show greater divergence in both sequence identity and structure (Fig. 2a, **Extended Data Fig. 9**). A comparison of the chromosome 5 *D5Z2* α-satellite HOR array, for example, reveals tracts that have as much as 4% sequence divergence, with clear expansions of α-satellite HORs in the CHM1 α-satellite HOR array (Fig. 2a).

**Figure 2.**
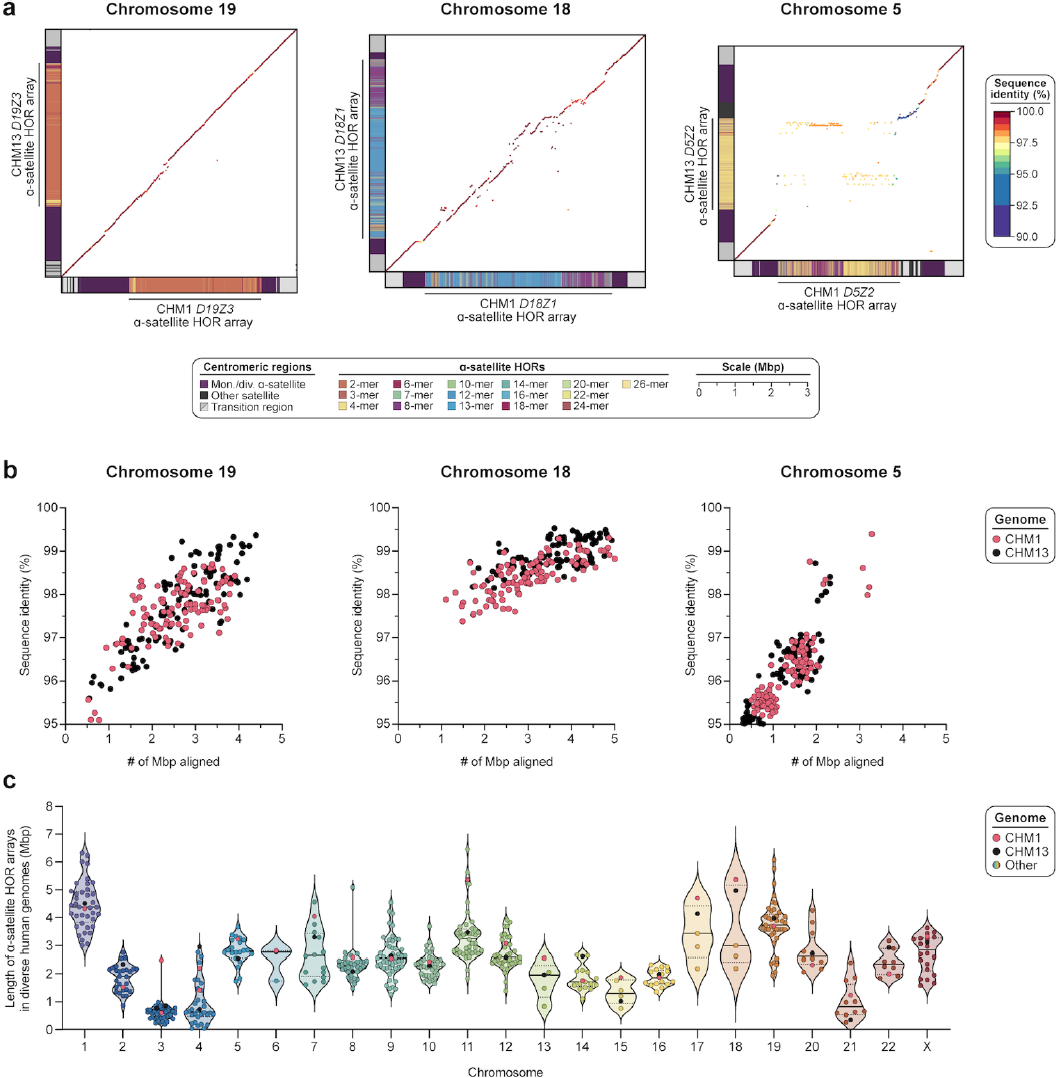
Variation in sequence and structure between two sets of human centromeres. **a)** Dot matrix plots showing allelic variation between CHM1 and CHM13 centromeric/pericentromeric haplotypes. Diagonal lines are colored by % sequence identity. The α-satellite HOR structure is shown on the axes, along with the organization of each centromeric/pericentromeric region. **b)** Comparison of the % sequence identity and # of Mbp aligned for 112 human centromere haplotypes from the HPRC^7^ and HGSVC^18^ mapped to the complete CHM1 and CHM13 centromere assemblies. Note that each dot represents a haplotype with 1:1 best mapping, although many of the centromeres are not yet complete in the HPRC/HGSVC samples. **c)** Plot showing the length of the active α-satellite HOR arrays among the CHM1 (red), CHM13 (black), and complete HPRC/HGSVC centromeres (various colors); n = 626. The α-satellite HOR arrays range in size from 0.03 Mbp on chromosome 4 to 6.5 Mbp on chromosome 11. Mean, solid black bar; 25% and 75% quartiles, dotted black bars.

Comparison with 56 incompletely assembled HPRC/HGSVC reference genomes^7^ generally confirms that this wide variance in sequence identity is a chromosome-specific property (Fig. 2b, **Extended Data Fig. 10**). While most α-satellite HOR arrays share at least 97% sequence identity, chromosomes 1, 5, 10, 12, 13, and 19 represent clear outliers, with 16.6% of α-satellite HOR arrays aligning very divergently (<97% sequence identity; **Extended Data Fig. 10**). Importantly, neither set of fully resolved human centromeres is a better match for the majority of HPRC/HGSVC genomes, nor does either adequately capture the full extent of human genetic diversity (**Extended Data Fig. 11**). For example, the mean sequence identity among the 56 HPRC/HGSVC genomes to either CHM1 or CHM13 is 98.0 ± 2.3% (**Extended Data Table 6**). Similarly, we find that 11 centromeres are a better match to CHM1, while 12 are a better match to CHM13 (**Extended Data Table 2**). If we require, however, that >75% of all HPRC haplotypes match better to either CHM1 or CHM13, only five centromeres meet this requirement for CHM1 (chromosomes 2, 12, 13, 19, and 22), while seven do for CHM13 (chromosomes 3, 4, 7, 10, 11, 14, and 15; **Extended Data Table 2**). These analyses reflect an extraordinary degree of single-nucleotide and structural diversity of human centromeres.

Comparison of the length of the α-satellite HOR arrays reveals that CHM1 arrays are ∼1.3-fold larger, on average, than their CHM13 counterparts, with 16 out of 23 chromosomes harboring a larger array in CHM1 than in CHM13 (Figs. 2c,3a, **Extended Data Table 7**). Of these, five arrays are >1.5-fold larger in CHM1 than in CHM13 (chromosomes 3, 4, 11, 15, and 21), with the greatest variation in length occurring on chromosome 21 (3.6-fold; Fig. 3a). This variation between CHM1 and CHM13 α-satellite HOR arrays falls within the normal range of variation (1.7- to 79.7-fold; median 2.3-fold), based on released haplotype-phased genome assemblies from the HPRC^7^ and HGSVC^18^ (Fig. 2c). Our analysis shows, for example, that human α-satellite HOR arrays range in size from 0.03 Mbp on chromosome 4 to 6.5 Mbp on chromosome 11. Chromosomes 3, 4, and 21 represent some of the smallest α-satellite HOR arrays as well as show the greatest variation in length among human haplotypes (Fig. 2c; 13.6-, 19.0-, 79.7-fold difference, respectively). Almost all the large-scale structural variation is due to variation in α-satellite HOR array organization and size, although the patterns are significantly more complex than simple insertion, deletion, or inversion processes (see below).

**Figure 3.**
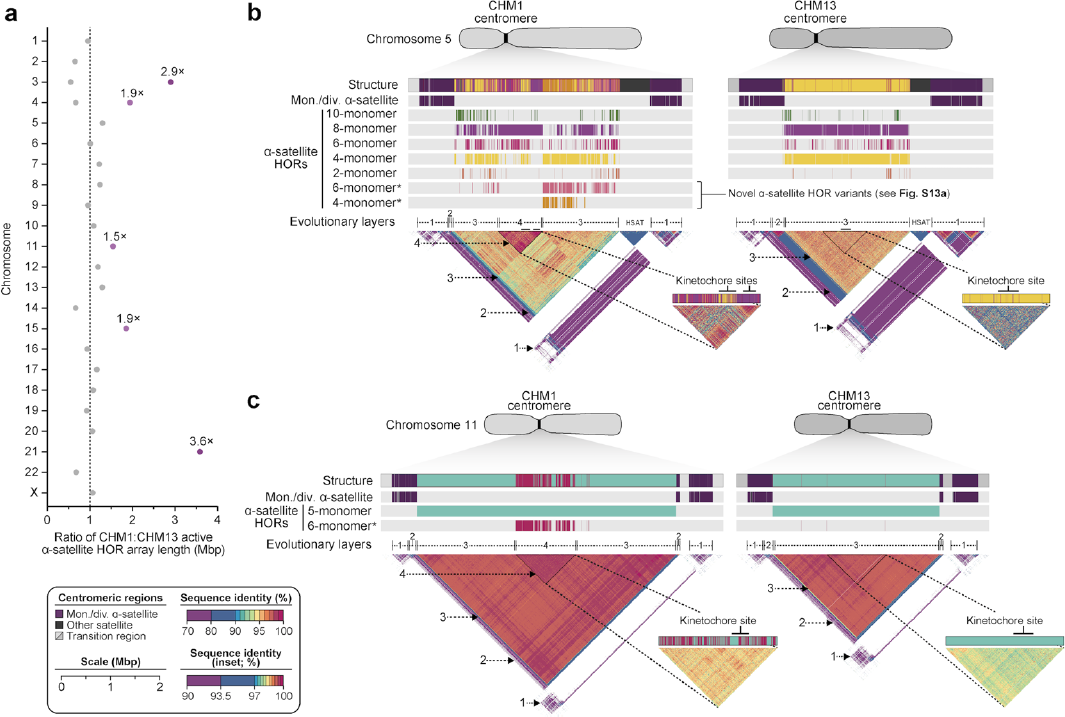
Variation in length and sequence composition of human centromeric α-satellite HOR arrays. **a)** Ratio of the length of the active α-satellite HOR arrays in the CHM1 genome compared to those in the CHM13 genome. **b,c)** Comparison of the **b)** CHM1 and CHM13 chromosome 5 *D5Z2* α-satellite HOR arrays and **c)** CHM1 and CHM13 chromosome 11 *D11Z1* α-satellite HOR arrays. The CHM1 chromosome 5 *D5Z2* array contains two novel α-satellite HOR variants as well as a new evolutionary layer (Layer 4; indicated with an arrow), which is absent from the CHM13 array. Similarly, the CHM1 chromosome 11 *D11Z1* α-satellite HOR array contains a 6-monomer HOR variant that is much more abundant than in the CHM13 array and comprises a new evolutionary layer (Layer 4; indicated with an arrow), although this 1.21-Mbp segment is more highly identical to the flanking sequence. The inset shows each of the new evolutionary layers with a higher stringency of sequence identity, as well as the relative position of the kinetochore.

Comparison of the α-satellite HOR arrays from the CHM1 and CHM13 centromeres identifies eight with distinctly different structures (chromosomes 5, 7, 8, and 10-14; Figs. 3b,c, **Extended Data Fig. 12**). This includes four arrays with a high abundance of previously uncharacterized α-satellite HORs (chromosomes 5, 7, 10, and 14; **Extended Data Fig. 13, Extended Data Table 8**). The centromeric *D5Z2* α-satellite array from CHM1 chromosome 5, for example, is significantly more diverse, containing two novel α-satellite HOR variants that are four and six α-satellite monomers in length (Fig. 3b, **Extended Data Fig. 13a**). Phylogenetic and comparative analysis of these HOR variants reveals that they are both derivatives of an ancestral 10-monomer α-satellite HOR, which resides at the edge of the *D5Z2* α-satellite HOR array. These novel HORs, confirmed by analysis of the HPRC genomes^7^, likely arose from repeated deletions of α-satellite monomers in the ancestral HOR, giving rise to novel 4- and 6-monomer HOR variants that subsequently propagated (**Extended Data Fig. 3a**). In addition, specific α-satellite HORs appear more consolidated, forming distinct layers that are not as apparent or are completely absent in the other haplotype. A clear 870-kbp layer, for example, is apparent in the CHM1 chromosome 5 centromeric *D5Z2* α-satellite HOR array, and it corresponds to a cluster of highly identical 8-monomer α-satellite HORs (Fig. 3b). This evolutionary layer is absent from the CHM13 centromere, whose 8-monomer α-satellite HORs are more dispersed along with the 4-monomer HORs. Similarly, the CHM1 chromosome 11 *D11Z1* centromere evolved a 1.2-Mbp layer in the core of its α-satellite HOR array that is missing from the CHM13 centromere (Fig. 3c). This novel layer is composed of 6-monomer α-satellite HORs that are found only rarely in the CHM13 centromere. We observe new evolutionary layers in the CHM1 chromosome 10, 12, and 13 α-satellite HOR arrays, all of which have divergent α-satellite HOR array structures. The remaining centromeres have a similar number of evolutionary layers between the two genomes, ranging from two to six, with the majority having four (**Extended Data Fig. 14**).

### Epigenetic differences between CHM1 and CHM13 centromeres

The kinetochore is a proteinaceous complex marked by the presence of nucleosomes containing the histone H3 variant CENP-A, which is critical to both meiotic and mitotic segregation of chromosomes. Prior studies have shown that the kinetochore typically resides within a region of hypomethylated DNA, dubbed the centromere dip region (CDR)^4, 26^, that colocalizes with CENP-A immunostaining^16^. We assessed the DNA methylation pattern and CENP-A chromatin organization of each CHM1 centromere and compared it to its CHM13 counterpart. Although CHM1 centromeric α-satellite HOR arrays are typically larger, the majority of CHM1 kinetochore sites (18 out of 23) are smaller than their CHM13 counterparts, with an average size of 178 versus 214 kbp, respectively (Fig. 4a, **Extended Data Table 7**). Additionally, 16 out of 23 CHM1 kinetochore sites are located more than 100 kbp away from their corresponding location in the CHM13 centromere, with six located more than 500 kbp away (chromosomes 4, 6, 11, 12, 18, and 20), when measuring the distance from the α-satellite HOR-to-monomeric transition region (Fig. 4b, **Extended Data Table 7**). Consistent with earlier observations^4^, we identify five chromosomes with evidence of two kinetochores, separated by >150 kbp (chromosomes 1, 2, 13, 17, and 19). In the case of chromosomes 13 and 19, the two distinct kinetochores are located more than 1 Mbp apart from each other (Fig. 4c,d). Assessment of the underlying sequence and structure of the chromosome 13 *D13Z2* α-satellite HOR array reveals a 631-kbp deletion in approximately half of CHM1 cells (**Extended Data Fig. 4, Supplementary Note 2**), suggesting that the two kinetochore sites likely represent two distinct cell populations or, alternatively, an early-stage somatic mutational event resulting in two differing genetic and epigenetic landscapes.

**Figure 4.**
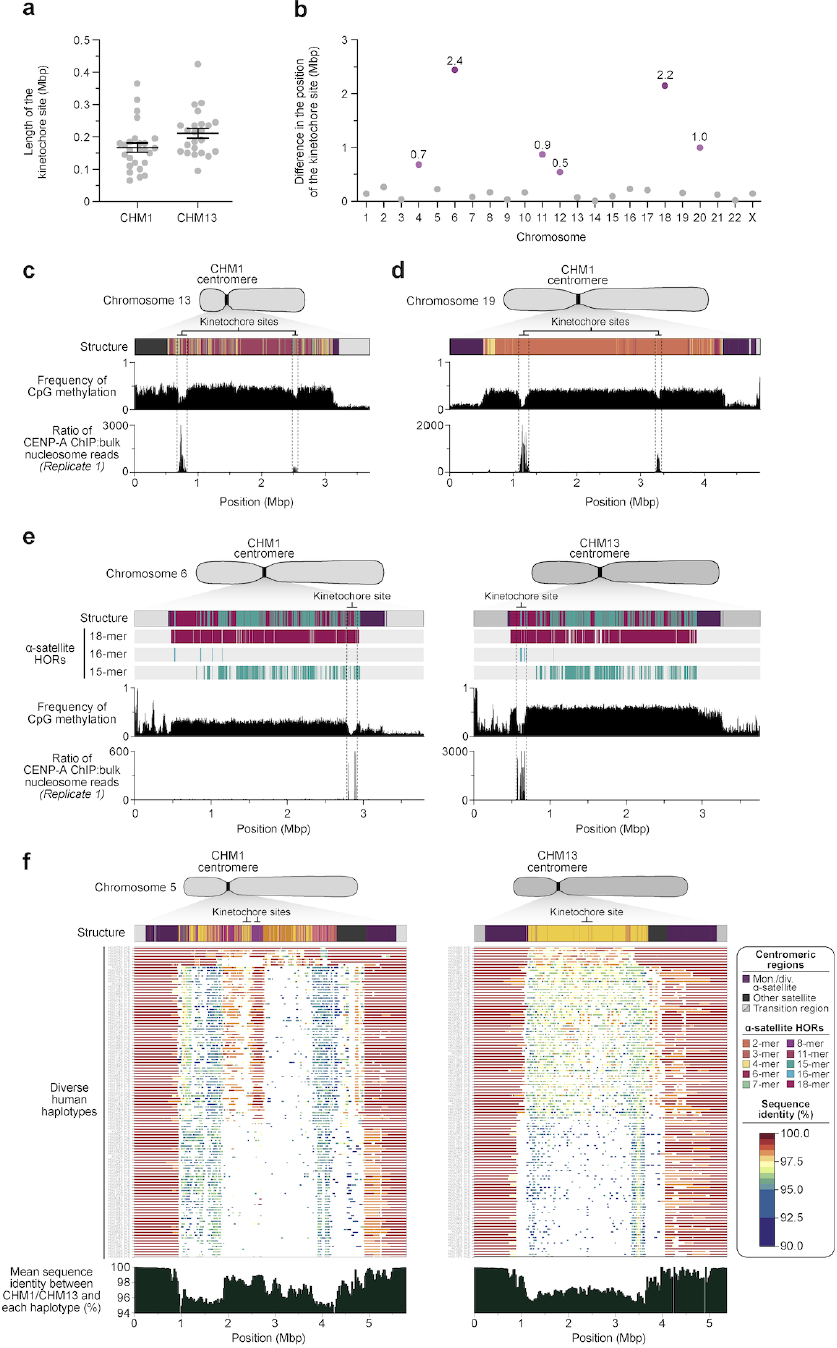
Variation in the site of the kinetochore among two sets of human centromeres. **a)** Plot comparing the length of the kinetochore site, marked by hypomethylated DNA and CENP-A-containing chromatin, between the CHM1 and CHM13 centromeres. **b)** Plot showing the difference in the position of the kinetochore among the CHM1 and CHM13 centromeres. **c,d)** Discovery of two potential kinetochores on the **c)** chromosome 13 and **d)** chromosome 19 centromeres in the CHM1 genomes. The presence of two hypomethylated regions enriched with CENP-A chromatin likely represents two populations of cells, which may have arisen due to a somatic mutation, resulting in differing epigenetic landscapes. **e)** Comparison of the CHM1 and CHM13 chromosome 6 centromeres, which differ in kinetochore position by 2.4 Mbp. **f)** Comparison of the CHM1 and CHM13 chromosome 5 centromeres, showing that the sequences underlying the CHM1 kinetochore are conserved in approximately half of the HPRC genomes, but the same degree of conservation is not observed for the CHM13 kinetochore region.

The chromosome 6 centromere shows the greatest variation in kinetochore position, with a difference of 2.4 Mbp between the two haplotypes. This change spans 87-88% of the length of the α-satellite HOR array itself and coincides with an alteration in the underlying α-satellite HOR sequence and structure, switching from a mixture of 16- and 18-monomer α-satellite HORs to a mixture of 15- and 18-monomer HORs (Fig. 4e). Given the complete sequence of CHM1 and CHM13 centromeres and the availability of incomplete assemblies from 56 diverse human genomes, we assessed whether the sequences underlying the kinetochore were more likely to be conserved compared to α-satellite HORs that were not associated with the kinetochore. While we observed clear examples of sequence conservation underlying the kinetochore for specific chromosomes involving both CHM1 and CHM13 (e.g., chromosomes 4, 5, 7, 12, 13, 16, and 18; Fig. 4f, **Extended Data Fig. 11**), other kinetochore regions appeared more similar (chromosomes 1-3, 6, 8, 9, 17, 20, 21, and X) or more divergent (chromosomes 10, 11, 14, 15, 19, and 22) than other portions of the α-satellite HOR array (**Extended Data Fig. 11**).

### Different centromere evolutionary trajectories among primate lineages

Our analyses (**Figs. 1-4**) revealed that human centromeres vary non-uniformly depending on the chromosome. In particular, specific human chromosomes show either highly variable α-satellite HOR array lengths (e.g., chromosome 21), diverse α-satellite HOR organizations (e.g., chromosomes 5, 10, and 12), or divergent epigenetic landscapes (e.g., chromosome 20). In contrast, the X chromosome is among the most conserved, with nearly identical sequences and structures among diverse human genomes (**Extended Data Fig. 11**). These findings imply that centromeres may have different mutation rates and diverse evolutionary trajectories that shape their variation. To test this hypothesis, we sequenced and assembled orthologous centromeres from four primate species, focusing on the completion of six centromeres, in an effort to reconstruct their evolutionary history over a 25-million-year window of primate evolution. To this end, we generated PacBio HiFi data (38- to 100-fold coverage) from diploid human, chimpanzee, orangutan, and macaque genomes (**Methods**), producing whole-genome assemblies ranging from 6.1 to 6.3 Mbp in size (**Extended Data Table 1**). Using ultra-long ONT data (14- to 20-fold coverage), we then ordered, oriented, and joined the PacBio HiFi contigs together from each centromere, creating 43 contiguous assemblies of primate centromeres for these six chromosomes (Fig. 5). Mapping of long-read sequencing data to each centromere showed uniform coverage, indicating a lack of large structural errors and validating the overall organization (**Extended Data Figs. 15,16**). With the exception of the X chromosome from a male chimpanzee, both haplotypes were completely sequenced for each diploid female sample, providing additional insights into their overall organization and variation (Fig. 5).

**Figure 5.**
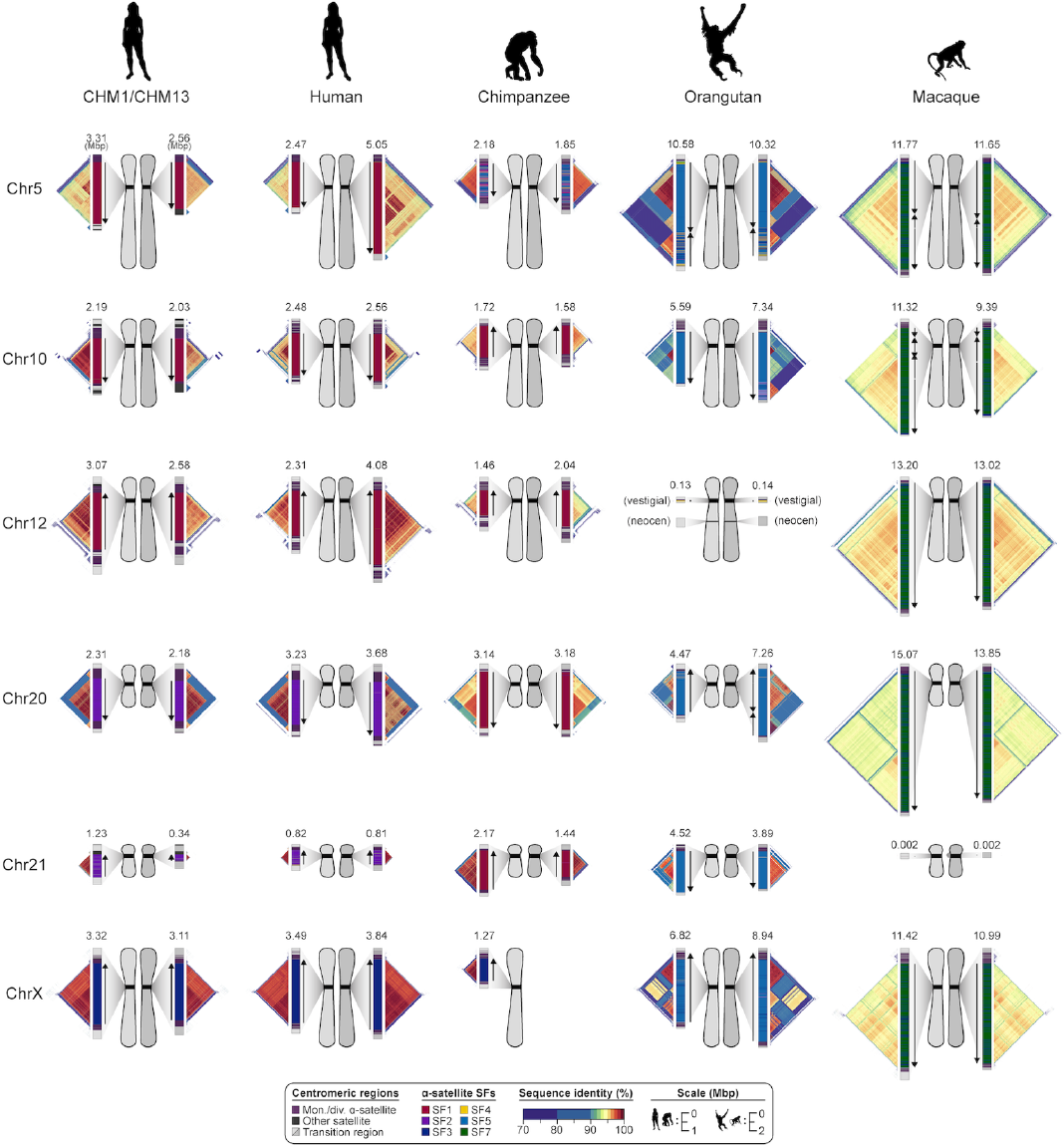
Sequence and structure of six sets of centromeres from diverse primate species. Complete assembly of centromeres from chromosomes 5, 10, 12, 20, 21, and X in human, chimpanzee, orangutan, and macaque reveals diverse α-satellite HOR organization and evolutionary landscapes. Sequence identity maps generated via StainedGlass^55^ are shown for each centromere (**Methods**), with the size of the α-satellite higher-order (human, chimpanzee, and orangutan) or dimeric (macaque) repeat array indicated in Mbp. The α-satellite suprachromosomal family (SF) for each centromeric array is indicated (vertical bar color), with arrows illustrating the orientation of the repeats within the array. Chromosome 12 in orangutan has a neocentromere, while the chromosome 21 centromere in macaque is no longer active due to a chromosomal fusion in that lineage^70^. All chromosomes are labeled according to the human phylogenetic group nomenclature^71^. The human diploid genome used as a control (second column) is HG00733—a 1000 Genomes sample of Puerto Rican origin. We note that the orangutan and macaque centromeres are drawn at half the scale with respect to the other apes.

Comparative analysis of these six sets of NHP centromeres revealed, as expected^13, 14, 27, 28^, diverse α-satellite HOR organization and structures, with α-satellite HOR arrays varying in size by more than 18.6-fold (the smallest residing on human chromosome 21, and the largest residing on macaque chromosome 20). Distinct species-specific differences also became apparent during this analysis (Fig. 5). For example, we estimate that common chimpanzee α-satellite HOR arrays are, on average, 67.8% the size of their human counterparts—a reduction observed in both chimpanzee haplotypes. Like humans, chimpanzee α-satellite HOR arrays show evidence of clear evolutionary layers, with the pairwise sequence identity of these layers dropping as they move toward pericentromeric DNA. This layered α-satellite HOR organization consists mainly of a single, continuous block of higher-order α-satellite repeats that are >95% identical to each other, except for on chromosomes 12 and 20, which have two or three discrete blocks of higher-order α-satellite repeats that are only 90-95% identical to each other. In contrast, orangutan centromere organization differs radically from either human or chimpanzee. We find that orangutan α-satellite HOR arrays are composed of three to four distinct blocks of α-satellite HORs that are only 80-90% identical to each other—thus, a mosaic of independent HOR expansions creating a “patchwork quilt” pattern based on sequence identity (Fig. 5). Finally, macaque centromeric α-satellite arrays are significantly larger in size, with an average length of 12.2 ± 1.6 Mbp. Unlike apes, which possess complex HOR structures, macaque centromeric arrays are composed of dimeric α-satellite units that are 93-97% identical across all centromeres.

Assessment of the suprachromosomal family (SF) relationships among each primate centromere revealed four unexpected findings. First, we identified the first African ape centromere that is primarily composed of SF5 α-satellite repeats: the chimpanzee chromosome 5 centromere. While all human and chimpanzee α-satellite arrays are mainly composed of α-satellites from SFs 1-3, we find that both chimpanzee chromosome 5 centromeres harbor a 2.1- or 2.4-Mbp region composed of SF5 α-satellite. This is exceptional, as the only other chimpanzee centromere known to be composed of non-SF 1-3 α-satellite is chromosome Y (SF4)^29^. Second, we found that all four chimpanzee chromosome 20 and 21 α-satellite HOR arrays are composed of SF1 α-satellite, as opposed to SF2 as in the human counterparts. SF1 is thought to be the ancestral version, as remnants of the SF2 arrays are found on the edges of the α-satellite HOR arrays. Third, we found that one orangutan chromosome 20 centromere harbors a large 3.2 Mbp inversion, while its homolog does not. This is the first polymorphic inversion found within an α-satellite HOR array within a single individual. Fourth, we found that all four macaque chromosome 5 and 10 α-satellite arrays harbor at least one large inversion—in the case of chromosome 5, both homologs have a 1.0- or 1.6-Mbp inversion; in the case of chromosome 10, only one homolog harbors a single inversion (2.8 Mbp), while the other homolog harbors two such inversions (1.6 and 2.5 Mbp) separated by a small (0.4 Mbp) stretch of directly oriented α-satellite dimers.

Despite these species-specific patterns, a common feature of all primate centromeres is the presence of two to five distinct evolutionary layers, marked by the most highly identical α-satellite sequences at the center of the satellite array that become increasingly divergent towards the periphery. These more divergent higher-order and dimeric repeats are flanked by blocks of monomeric α-satellite DNA. We performed phylogenetic and comparative analyses of all six complete orthologous centromere sets and observed that monomeric α-satellite is generally more closely related to the Old World monkey dimeric satellites of macaques. Notwithstanding this general topology, distinct chromosome-specific patterns emerge (Fig. 6, **Extended Data Fig. 17**). The chromosome 5 centromere, for example, has evolved human-specific α-satellite that define the active *D5Z2* α-satellite HOR array, while more ancient α-satellite sequences are located within inactive *D5Z1* α-satellite HOR arrays (Fig. 6a). This is in contrast to the chromosome 12 centromere, which harbors α-satellite HORs that are shared among orangutan and chimpanzee (Fig. 6b). Finally, the chromosome X centromere is composed of α-satellite HORs and monomers that are evolutionarily similar to each other, and unlike the other centromeres, are also similar to macaque’s α-satellite monomers (Fig. 6c).

**Figure 6.**
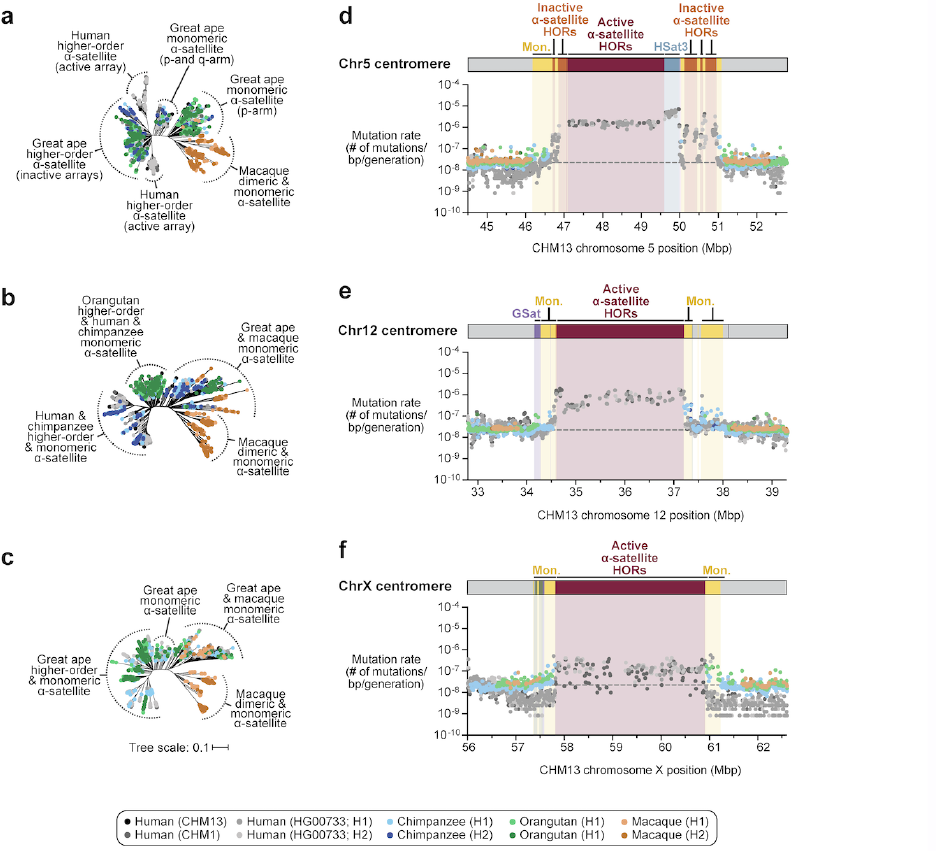
Centromeres evolve with different evolutionary trajectories and mutation rates. **a-c)** Phylogenetic trees of human, chimpanzee, orangutan, and macaque α-satellites from the higher-order and monomeric α-satellite regions of the chromosome 5, 12, and X centromeres, respectively. **d-f)** Plot showing the mutation rate of the chromosome 5, 12, and X centromeric regions, respectively. Individual data points from 10 kbp pairwise sequence alignments are shown.

### Human haplotype phylogenetic reconstruction and mutation rate estimates

Because our analyses showed that the monomeric α-satellite sequences mutate less quickly and can be readily aligned among human and nonhuman apes, we focused first on the pericentromeric DNA flanking the α-satellite HOR array. Based on complete sequence from human and NHP centromeric transition regions, we estimated the mutation rate of the ∼1-2 Mbp region flanking the α-satellite HOR arrays using established evolutionary models (**Methods**) and found the mutation rate increases 1.1- to 4.1-fold compared to the unique portions in each of the six centromeres (Fig. 6d-f, **Extended Data Fig. 17d-f**). The greatest increase in mutation rate is observed for the chromosome 5 centromere (4.1-fold), while the smallest increase occurs for the chromosome X centromere (1.1-fold), consistent with the observed rapid and slower structural diversity for this chromosome. Due to nearly complete evolutionary turnover of the α-satellite HORs, biologically meaningful alignment comparisons among humans and nonhuman apes could not be made. However, analyses of the sequence alignments among the four human haplotypes suggested a potential mutation rate increase of at least an order of magnitude, given the caveat that significant portions of the α-satellite repeats failed to align.

To understand the nature of evolutionary change within the α-satellite HOR arrays and especially the emergence of new HORs, we applied a population genetics approach leveraging the genetic diversity present in the HPRC^7^ and HGSVC^18^ genomes. We reasoned that less divergent sequence comparisons within the human species would allow for more accurate alignments and, therefore, better reconstruction of the series of mutational events occurring within the α-satellite HOR arrays. Given the relative stability of the flanking monomeric satellite DNA, we constructed phylogenetic trees using the chimpanzee sequence as an outgroup and estimated separation times for different human haplotypes, assuming a chimpanzee and human divergence time of 6 million years (Fig. 7a, **Extended Data Figs. 18-20**). Under the assumption that there is limited or no recombination across the α-satellite HOR array, we then compared the topologies of both the p- and q-arms, focusing specifically on haplotypes where we had documented the emergence of novel α-satellite HOR arrays. Despite being anchored in sequence separated 2-3 Mbp apart, the p- and q-arm topologies of the resulting trees were remarkably similar, consistent with the notion of suppressed or limited recombination across the region. Importantly, haplotypes harboring new α-satellite HORs most often share a monophyletic origin (Fig. 7b,c, **Extended Data Figs. 18-20**). For example, in the case of chromosome 12, we estimate the new HORs emerged approximately 13-23 kya (thousand years ago; Fig. 7b), while for chromosome 11, they emerged approximately 80-153 kya (Fig. 7c). This suggests a single origin for the new α-satellite HORs, followed by the saltatory spread of >1 Mbp of new HORs to this subset of human haplotypes.

**Figure 7.**
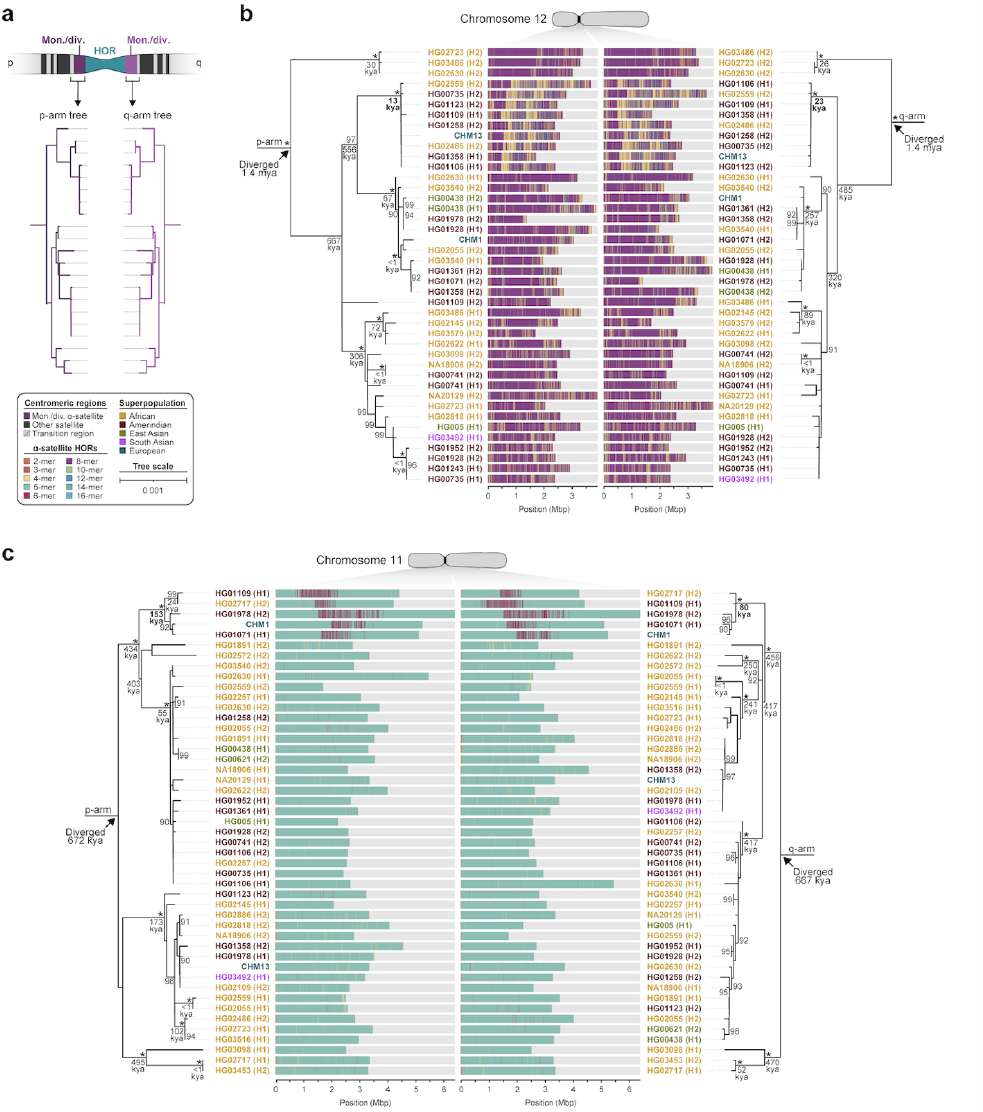
Phylogenetic reconstruction of human centromeric haplotypes and the saltatory amplification of new α-satellite HORs. **a)** Strategy to determine the phylogeny and divergence times of completely sequenced centromeres using monomeric α-satellite or unique sequence flanking the canonical α-satellite HOR array from both the short (p) and long (q) arms of chromosomes 11 and 12. Chimpanzee is used as an outgroup with an estimated species divergence time of 6 million years ago. **b,c)** Maximum-likelihood phylogenetic trees depicting the p- and q-arm topologies along with the estimated divergence times reveals a monophyletic origin for the emergence of new α-satellite HORs within the **b)** chromosome 12 (*D12Z3*) and **c)** chromosome 11 (*D11Z1*) α-satellite HOR arrays. These arrays show a complex pattern of new α-satellite HOR insertions and deletions over a short period of evolutionary time.

By directly comparing the structure of the α-satellite HOR arrays with the nearest human haplotype lacking the newly derived HORs, we computed the difference in the number of base pairs, α-satellite monomer units, α-satellite HOR units, and distinct structural changes (**Extended Data Table 9**). Using these newly minted α-satellite HORs as a benchmark, our results suggest 392-2,490 nucleotide differences (or up to two α-satellite HORs) per generation, on average, to create the new HORs on chromosomes 11 and 12 (Fig. 7b,c). Given the average length of each α-satellite HOR array and the estimated coalescent time, this translates to remarkably different rates for the emergence of these new α-satellite HORs on chromosomes 11 (∼30-60 nucleotide differences per Mbp per generation) and 12 (∼500-1000 nucleotide differences per Mbp per generation; **Extended Data Table 9**). While caution should be exercised given the focus on new α-satellite HOR structures and the limited number of human haplotypes compared, a surprising finding is both the speed at which these new HORs emerged and the interdigitated nature of new α-satellite HORs intermixed with relic ancestral HORs. Our results suggest approximately 100 distinct structural changes (insertions and deletions) as this new HOR variant evolved. This pattern implicates mechanisms other than simple unequal crossover for the spread of novel α-satellite HORs within centromeres. The incredible change in array structure is likely due to saltatory amplification of newly emerged α-satellite HOR variants at multiple sites in the original HOR array, leading to an overall increase in array size from 554 kbp to 2 Mbp, on average (Fig. 7b,c, **Extended Data Figs. 18-20**).

## DISCUSSION

We present a detailed comparative analysis of two completely assembled reference sets of human centromeres compared to a diversity panel of human and NHP centromeres. We show a demonstrable acceleration of single-nucleotide and structural variation transitioning from euchromatin to heterochromatin, with most of this excess occurring within the core of the centromeric α-satellite HOR arrays. Consequently, active α-satellite HOR arrays vary substantially in size and structure, with the smallest arrays residing on chromosomes 3, 4, and 21 (Fig. 2c; 0.2, 0.03, and 0.3 Mbp, respectively), and the largest arrays residing on chromosomes 1, 11, 17, and 18 (Fig. 2c; 6.3, 6.5, 4.7, and 5.4 Mbp, respectively). There are, however, two important caveats to the current analysis. First, the length and variance of these arrays are based on those centromeres that are contiguously and accurately assembled, creating a potential ascertainment bias if smaller centromeres preferentially assemble. Second, a significant fraction (45-47%) of the completely sequenced centromeres cannot be readily aligned to either of the two references due, in part, to the emergence of new α-satellite HOR structures (**Extended Data Table 6**). Focusing just on the two human haplotypes represented by CHM1 and CHM13, we find no less than eight chromosomes that have distinctly different α-satellite HOR array structures (chromosomes 5, 7, 8, and 10-14; Fig. 3b,c, **Extended Data Fig. 12**) and four that harbor novel α-satellite HOR variants in high abundance (chromosomes 5, 7, 10, and 14; Fig. 3b, **Extended Data Fig. 13**, **Extended Data Table 8**). Our estimates of genetic diversity and mutation rate, thus, likely represent an underestimate until a greater diversity of human haplotypes, including large and more structurally diverse centromeres, are fully sequence resolved.

Interestingly, the site of the kinetochore attachment (marked by hypomethylated DNA and an enrichment of chromatin containing the centromeric histone H3 variant, CENP-A) varies considerably between the two human reference sets. We found that approximately two-thirds of CHM1 centromeres have a kinetochore located at least 100 kbp away from their corresponding position in the CHM13 reference genome (Fig. 4b). Eight, in fact, differ by more than 500 kbp (chromosomes 5, 6, 11-13, and 18-20; Fig. 4b), with a few showing evidence of more than one location in CHM1 (Fig. 4c,d). Some centromeres (chromosomes 5 and 11) show that the repositioning of the kinetochore corresponds with the emergence of a novel evolutionary layer within the core of the α-satellite HOR array (Fig. 3b,c). We hypothesized that the kinetochore may favor particular DNA sequences or motifs that differ from the remainder of the α-satellite HOR array. However, comparison of the α-satellite sequences enriched with, or devoid of, CENP-A from each of the CHM1 and CHM13 centromeres failed to reveal a clear association (Fig. 4f; **Extended Data Fig. 11**). This remarkable plasticity in kinetochore position despite the conserved, essential function of these regions underscores the “centromere paradox”^30^, an unresolved conundrum regarding the contradictory phenomenon of rapidly evolving centromeric DNA and proteins despite their essential role in ensuring faithful chromosome transmission. The germline and somatic stability of both the kinetochore location and the underlying DNA sequence will need to be investigated by examining genetic and epigenetic variation in centromeres both across generations and in multiple primary tissues from the same donor.

The considerable variability in sequence, structure, and epigenetic landscape among human centromeres led us to hypothesize that centromeres may mutate at different rates and with different evolutionary trajectories. To test this hypothesis, we sequenced and assembled six sets of orthologous centromeres from the chimpanzee, orangutan, and macaque genomes, resolving both haplotypes for each centromere (except for the chromosome X in the male chimpanzee), for a total of 31 fully resolved centromeres from NHPs (Fig. 5). Comparison of the sequence and structure across each primate centromere revealed unique features specific to their α-satellite arrays. We find that chimpanzee α-satellite HOR arrays tend to be 67-68% smaller than the human counterparts with some peculiar idiosyncrasies. The chimpanzee chromosome 5 centromere, for example, is primarily composed of SF5 α-satellite repeats, unlike all other human and chimpanzee centromeres (except for chromosome Y in both species), which are composed of SF1-3 α-satellite repeats. Orangutan centromeres tend to have a single α-satellite HOR array, but unlike other apes, the array is composed of a mosaic “patchwork” of distinct α-satellite HOR blocks with a high degree of divergence. Macaque centromeres are consistently the largest, but also more homogenous, composed of dimeric α-satellites that are approximately 95% similar in sequence to each other. Macaque also has the distinction of harboring novel polymorphic inversions in its dimeric array (e.g., chromosome 5 and 10 centromeres).

Comparative and phylogenetic analysis of the α-satellite monomers from each primate centromere suggests near-complete species-specific turnover of the α-satellite HOR structures among primate species. Analyses of variation among human haplotypes, however, shows that each centromere mutates at a different rate. The chromosome 5 centromere, for example, mutates at least 10-fold faster than the chromosome X centromere, with the net effect that almost 48% of the α-satellite HORs cannot be aligned to either CHM1 or CHM13 references (Fig. 6). This rapid evolution has led to the emergence of new, human-specific α-satellite HORs that are unique to a subset of haplotypes. Using the emergence of these new HOR structures within human as a marker of evolutionary mutability, we developed an approach to estimate the rate of evolutionary change by comparing closely related finished centromeres within a coalescent framework. Our results for two human centromeres (chromosomes 11 and 12; Fig. 7) suggest that centromeric α-satellite HOR arrays can mutate multiple orders of magnitude more quickly than unique DNA (estimated at 30-1000 nucleotides per generation per Mbp based on our analysis of newly emerged HORs on chromosomes 11 and 12; **Extended Data Table 9**). These changes in DNA occur most frequently in concert with gains and losses of α-satellite HOR units and do not appear to do so in a contiguous manner but, instead, are intermixed with ancestral HORs. The mechanism responsible for these changes is currently not well described, but it is hypothesized that they occur in a saltatory fashion as opposed to a constant rate of mutation^31^, potentially as a result of meiotic drive for the newly minted HORs^32^. The expansion in the length of the α-satellite arrays upon the emergence of new α-satellite HORs may also contribute to increased centromere strength^33^, which can lead to non-Mendelian chromosome segregation and biased chromosome retention in oocytes^34, 35^. Now that centromeres can be fully phased and assembled, it will be critical to study the mutational processes in a multigenerational families to understand the mechanisms shaping these rapidly evolving regions of our genome. Uncovering one of these new α-satellite HOR variants transmitting within a family will likely provide critical insights into mechanisms underlying centromere mutation and evolution.

## MATERIALS AND METHODS

### Cell lines

CHM1hTERT (abbr. CHM1) cells were originally isolated from a hydatidiform mole at Magee-Womens Hospital (Pittsburgh, PA). Cryogenically frozen cells from this culture were grown and transformed using human telomerase reverse transcriptase (TERT) to immortalize the cell line. This cell line has been authenticated via STR analysis by Cell Line Genetics (Madison, WI) and has tested negative for mycoplasma contamination. Human HG00733 lymphoblastoid cells were originally obtained from a female Puerto Rican child, immortalized with the Epstein-Barr Virus (EBV), and stored at the Coriell Institute for Medical Research (Camden, NJ). Chimpanzee (*Pan troglodytes*; Clint; S006007) fibroblast cells were originally obtained from a male western chimpanzee named Clint (now deceased) at the Yerkes National Primate Research Center (Atlanta, GA) and immortalized with EBV. Orangutan (*Pongo abelii*; Susie; PR01109) fibroblast cells were originally obtained from a female Sumatran orangutan named Susie (now deceased) at the Gladys Porter Zoo (Brownsville, TX), immortalized with EBV, and stored at the Coriell Institute for Medical Research (Camden, NJ). Macaque (*Macaca mulatta*; AG07107) fibroblast cells were originally obtained from a female rhesus macaque of Indian origin and stored at the Coriell Institute for Medical Research (Camden, NJ). The HG00733, chimpanzee, orangutan, and macaque cell lines have not yet been authenticated or assessed for mycoplasma contamination to our knowledge.

### Cell culture

CHM1 cells were cultured in complete AmnioMax C-100 Basal Medium (Thermo Fisher Scientific, 17001082) supplemented with 15% AmnioMax C-100 Supplement (Thermo Fisher Scientific, 12556015) and 1% penicillin-streptomycin (Thermo Fisher Scientific, 15140122). HG00733 (*Homo sapiens*) cells were cultured in RPMI-1650 media (Sigma Aldrich, R8758) supplemented with 15% fetal bovine serum (FBS; Thermo Fisher Scientific, 16000-044) and 1% penicillin-streptomycin (Thermo Fisher Scientific, 15140122). Chimpanzee (*Pan troglodytes*; Clint; S006007) and macaque (*Macaque mulatta*; AG07107) cells were cultured in MEM α containing ribonucleosides, deoxyribonucleosides, and L-glutamine (Thermo Fisher Scientific, 12571063) supplemented with 12% FBS (Thermo Fisher Scientific, 16000-044) and 1% penicillin-streptomycin (Thermo Fisher Scientific, 15140122). Orangutan (*Pongo abelii*; Susie; PR01109) cells were cultured in MEM α containing ribonucleosides, deoxyribonucleosides, and L-glutamine (Thermo Fisher Scientific, 12571063) supplemented with 15% FBS (Thermo Fisher Scientific, 16000-044) and 1% penicillin-streptomycin (Thermo Fisher Scientific, 15140122). All cells were cultured in a humidity-controlled environment at 37°C with 95% O_2_.

### DNA extraction, library preparation, and sequencing

PacBio HiFi data were generated from the CHM1 and HG00733 genomes as previously described^16^, with some modifications. Briefly, high-molecular-weight (HMW) DNA was extracted from cells using a modified Qiagen Gentra Puregene Cell Kit protocol^36^. HMW DNA was used to generate PacBio HiFi libraries via the Template Prep Kit v1 (PacBio, 100-259-100) or SMRTbell Express Template Prep Kit v2 (PacBio, 100-938-900) and SMRTbell Enzyme Clean Up kits (PacBio, 101-746-400 and 101-932-600). Size selection was performed with SageELF (Sage Science, ELF001), and fractions sized 11, 14, 15, or 16 kbp [as determined by FEMTO Pulse (Agilent, M5330AA)] were chosen for sequencing. Libraries were sequenced on the Sequel II platform with seven or eight SMRT Cells 8M (PacBio, 101-389-001) per sample using either Sequel II Sequencing Chemistry 1.0 (PacBio, 101-717-200) or 2.0 (PacBio, 101-820-200), both with 2-hour pre-extension and 30-hour movies, aiming for a minimum estimated coverage of 30X in PacBio HiFi reads (assuming a genome size of 3.1 Gbp). Raw CHM1 data was processed with DeepConsensus^37^ (v0.2.0) with the default parameters. Raw HG00733 data was processed using the CCS algorithm (v3.4.1) with the following parameters: –minPasses 3 – minPredictedAccuracy 0.99 –maxLength 21000 or 50000.

Ultra-long ONT data were generated from the CHM1, HG00733, chimpanzee, orangutan, and macaque genomes according to a previously published protocol^38^. Briefly, 3-5 x 10^7^ cells were lysed in a buffer containing 10 mM Tris-Cl (pH 8.0), 0.1 M EDTA (pH 8.0), 0.5% w/v SDS, and 20 ug/mL RNase A (Qiagen, 19101) for 1 hour at 37°C. 200 ug/mL Proteinase K (Qiagen, 19131) was added, and the solution was incubated at 50°C for 2 hours. DNA was purified via two rounds of 25:24:1 phenol-chloroform-isoamyl alcohol extraction followed by ethanol precipitation. Precipitated DNA was solubilized in 10 mM Tris (pH 8.0) containing 0.02% Triton X-100 at 4°C for two days. Libraries were constructed using the Ultra-Long DNA Sequencing Kit (ONT, SQK-ULK001) with modifications to the manufacturer’s protocol. Specifically, ∼40 ug of DNA was mixed with FRA enzyme and FDB buffer as described in the protocol and incubated for 5 minutes at RT, followed by a 5-minute heat-inactivation at 75°C. RAP enzyme was mixed with the DNA solution and incubated at RT for 1 hour before the clean-up step. Clean-up was performed using the Nanobind UL Library Prep Kit (Circulomics, NB-900-601-01) and eluted in 225 uL EB. 75 uL of library was loaded onto a primed FLO-PRO002 R9.4.1 flow cell for sequencing on the PromethION, with two nuclease washes and reloads after 24 and 48 hours of sequencing.

Additional ONT data was generated from the CHM1, HG00733, chimpanzee, orangutan, and macaque genomes according to a previously published protocol^16^. Briefly, HMW DNA was extracted from cells using a modified Qiagen Gentra Puregene protocol^36^. HMW DNA was prepared into libraries with the Ligation Sequencing Kit (SQK-LSK110) from ONT and loaded onto primed FLO-PRO002 R9.4.1 flow cells for sequencing on the PromethION, with two nuclease washes and reloads after 24 and 48 hours of sequencing. All ONT data were base-called with Guppy (v5.0.11) with the SUP model.

### Targeted sequence assembly and validation of centromeric regions

To generate complete assemblies of centromeric regions from the CHM1, HG00733, chimpanzee, orangutan, and macaque genomes, we first assembled each genome from PacBio HiFi data (**Extended Data Table 1**) using hifiasm^19^ (v0.16.1). The resulting PacBio HiFi contigs were aligned to the T2T-CHM13 reference genome^3^ (v2.0) via minimap2^39^ (v2.24) with the following parameters: -I 15G -a --eqx-x asm20 -s 5000. Fragmented centromeric contigs were subsequently scaffolded with ultra-long (>100 kbp) ONT data generated from the same source genome via a method that takes advantage of SUNKs (**Extended Data Fig. 1**; https://github.com/arozanski97/SUNK-based-contig-scaffolding). Briefly, SUNKs (*k*=20 bp) were identified from the CHM1 PacBio HiFi whole-genome assembly via Jellyfish (v2.2.4) and barcoded on the CHM1 PacBio HiFi centromeric contigs as well as all ultra-long ONT reads. PacBio HiFi centromeric contigs sharing a SUNK barcode with ultra-long ONT reads were subsequently joined together to generate contiguous assemblies that traverse each centromeric region. The base accuracy of the assemblies was improved by replacing the ONT sequences with locally assembled PacBio HiFi contigs generated via HiCanu^2^ (v2.1.1).

We validated the construction of each centromere assembly with four different methods. First, we aligned native PacBio HiFi and ONT data from the same source genome to each whole-genome assembly using pbmm2 (v1.1.0) (for PacBio HiFi data; https://github.com/PacificBiosciences/pbmm2) or Winnowmap^40^ (v1.0) (for ONT data) and assessed the assemblies for uniform read depth across the centromeric regions via IGV^41^ and NucFreq^17^. Next, we assessed the concordance between the assemblies and raw PacBio HiFi data using VerityMap^22^, which identifies discordant *k*-mers between the two and flags them for correction. Then, we assessed the concordance between the assemblies and ONT data using GAVISUNK^23^, which identifies concordant SUNKs between the two. Finally, we estimated the accuracy of the centromere assemblies from mapped *k*-mers (*k*=21) using Merqury^42^ and publicly available Illumina data from each genome (**Extended Data Table 1**). We estimated the QV of the centromeric regions with the following formula:

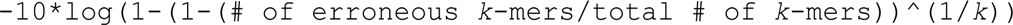

### Fluorescent *in situ* hybridization and spectral karyotyping

To determine the karyotype of the CHM1 genome, we first prepared metaphase chromosome spreads by arresting CHM1 cells in mitosis via the addition of KaryoMAX Colcemid Solution (0.1 µg/ml, Thermo Fisher Scientific, 15212012) to the growth medium for 6 hours. Cells were collected by centrifugation at 200*g* for 5 minutes and incubated in 0.4% KCl swelling solution for 10 min. Swollen cells were pre-fixed by the addition of freshly prepared methanol:acetic acid (3:1) fixative solution (∼100 μL per 10 ml total volume). Pre-fixed cells were collected by centrifugation at 200*g* for 5 min and fixed in methanol:acetic acid (3:1) fixative solution. Spreads were dropped on a glass slide and incubated on a heating block at 65°C overnight. Before hybridization, slides were treated with 1 mg/ml RNase A (1 mg/ml, Qiagen, 19101) in 2x SSC for at least 45 min at 37°C and then dehydrated in a 70%, 80%, and 100% ethanol series for 2 minutes. Denaturation of spreads was performed in 70% formamide/2X SSC solution at 72°C for 1.5 minutes and immediately stopped by immersing slides in an ethanol series pre-chilled to - 20°C.

Fluorescent probes for spectral karyotyping were generated in-house. Individual fluorescently labeled whole-chromosome paints were obtained from Applied Spectral Imaging. Paints were provided in a hybridization buffer and mixed 1:1 for indicated combinations. Labeled chromosome probes and paints were denatured by heating to 80°C for 10 minutes before applying them to denatured slides. Spreads were hybridized to probes under a HybriSlip hybridization cover (Grace Bio-Labs, 716024) sealed with Cytobond (SciGene, 2020-00-1) in a humidified chamber at 37°C for 48 hr. After hybridization, slides were washed in 50% formamide/2X SSC 3 times for 5 min at 45°C, 1x SSC solution at 45°C for 5 min twice, and at room temperature once. Slides were then rinsed with double-deionized H_2_O, air-dried, and mounted in Vectashield containing DAPI (Vector Laboratories, H-1200-10).

For spectral karyotyping, images were acquired using LSM710 confocal microscope (Zeiss) with the 63x/1.40 NA oil objective. Segmentation, spectral unmixing, and identification of chromosomes were performed using an open-source KISS (Karyotype Identification via Spectral Separation) analysis package for Fiji^43^, freely available at http://research.stowers.org/imagejplugins/KISS_analysis.html. For a detailed description of chromosome paints, hybridization, and analysis procedures, see Ref. 44.

For individually painted chromosomes, Z-stack images were acquired on the Nikon TiE microscope equipped with a 100x objective NA 1.45, Yokogawa CSU-W1 spinning disk, and Flash 4.0 sCMOS camera. Image processing was performed in Fiji^43^.

### Strand-seq analysis

To assess the karyotype of the CHM1 genome, we prepared Strand-seq libraries from CHM1 cells using a previously published protocol^45, 46^. We sequenced the mono- and di-nucleosome fractions separately, with the mononuclesomes sequenced with 75 bp, paired-end Illumina sequencing, and the dinucleosomes sequenced with 150 bp, paired-end Illumina sequencing. We demultiplexed the raw sequencing data based on library-specific barcodes and converted them to FASTQ files using Illumina standard software. We aligned the reads in the FASTQ files to the T2T-CHM13 reference genome^3^ (v2.0) using BWA^47^ (v0.7.17-r1188), sorted the alignments using SAMtools^48^ (v1.9), and marked duplicate reads with sambamba^49^ (v1.0). We merged the BAM files for the mono- and di-nucleosome fractions of each cell using SAMtools^48^ (v1.9). We used breakpointR^50^ to assess the quality of generated Strand-seq libraries with the following parameters: windowsize = 2000000, binMethod = ’size’, pairedEndReads = TRUE, min.mapq = 10, background = 0.1, minReads = 50. We filtered the libraries based on read density, level of background reads, and level of genome coverage variability^51^. A total of 48 BAM files were selected for all subsequent analysis and are publicly available. We detected changes in strand-state inheritance across all Strand-seq libraries using the R package AneuFinder^52^ with the following parameters: variable.width.reference = <MERGED BAM of all 48 Strand-seq libraries>, binsizes = windowsize, use.bamsignals = FALSE, pairedEndReads = TRUE, remove.duplicate.reads = TRUE, min.mapq = 10, method = ’edivisive’, strandseq = TRUE, cluster.plots = TRUE, refine.breakpoints = TRUE. We extracted a list of recurrent strand-states changes reported as sister chromatid exchange hotspots by AneuFinder. With this analysis, we identified reciprocal translocations between chromosomes 4q35.1/11q24.3 and 16q23.3/17q25.3 (see below) and established the overall copy number for each chromosome and Strand-seq library.

To identify the reciprocal translocation breakpoints between chromosomes 4q35.1/11q24.3 and 16q23.3/17q25.3 in the CHM1 genome, we first aligned CHM1 PacBio HiFi reads to the T2T-CHM13 reference genome^3^ (v2.0) via pbmm2 (v1.1.0) and used BEDtools^53^ intersect (v2.29.0) to define putative translocation regions based on AneuFinder analysis (described above). We extracted PacBio HiFi reads with supplementary alignments with SAMtools^48^ flag 2048. Through this method, we were able to identify the precise breakpoint of each translocation. We note that for the reciprocal translocation between chromosomes 4q35.1/11q24.3, we report two breakpoints in each chromosome due to the presence of a ∼97-98 kbp deletion in the translocated homologs (**Extended Data Fig. 3**). The breakpoints are located at chr4:187112496/chr11:130542388, chr4:187209555/chr11:130444240, and chr16:88757545/chr17:81572367 (in T2T-CHM13 v2.0).

### Sequence identity across centromeric regions

To calculate the sequence identity across the centromeric regions from CHM1, CHM13, and 56 other diverse human genomes (generated by the HPRC^7^ and HGSVC^18^), we performed three analyses that take advantage of different alignment methods. In the first analysis, we performed a pairwise sequence alignment between contigs from the CHM1, CHM13, and diverse genomes using minimap2^39^ (v2.24) and the following command: minimap2 -I 15G -K 8G -t {threads} -ax asm20 --secondary=no --eqx -s 2500 {ref.fasta} {query.fasta}. We filtered the alignments using SAMtools^48^ (v1.9) flag 4, which keeps primary and partial alignments. We subsequently partitioned the alignments into 10-kbp non-overlapping windows in the reference genome (either CHM1 or CHM13) and calculated the mean sequence identity between the pairwise alignments in each window. We averaged the sequence identity across the 10-kbp windows within the α-satellite higher-order repeat (HOR) array(s), monomeric/diverged α-satellites, other satellites, and non-satellites for each chromosome to determine the mean sequence identity in each region.

In the second analysis, we first fragmented the centromeric contigs from each genome assembly into 10-kbp fragments with seqtk (v1.3; https://github.com/lh3/seqtk) and subsequently aligned them to the reference genome (either CHM1 or CHM13) using minimap2^39^ (v2.24) and the following command: minimap2 -I 15G -K 8G -t {threads} -ax asm20 --secondary=no --eqx -s 40 {ref.fasta} {query.fasta}. We filtered the alignments using SAMtools^48^ (v1.9) flag 4, which keeps primary and partial alignments. In this method, multiple 10-kbp fragments are allowed to align to the same region in the reference genome, but each 10-kbp fragment is only allowed to align once. We, then, partitioned the alignments into 10-kbp non-overlapping windows in the reference genome and calculated the mean sequence identity between all alignments in each window. We averaged the sequence identity across the 10-kbp windows within the α-satellite HOR array(s), monomeric/diverged α-satellites, other satellites, and non-satellites for each chromosome to determine the mean sequence identity in each region.

In the third analysis, we first identified the location of the α-satellite HOR array(s) in each genome assembly using RepeatMasker^54^ (v4.1.0) followed by Hum-AS-HMMER (https://github.com/fedorrik/HumAS-HMMER_for_AnVIL) and subsequently extracted regions enriched with “live” α-satellite HORs (denoted with an “L” in the Hum-AS-HMMER BED file). We, then, ran TandemAligner^25^t (v0.1) on pairs of complete centromeric HOR arrays using the following command: tandem_aligner --first {ref.fasta} --second {query.fasta} -o {output_directory}. We parsed the CIGAR string generated by TandemAligner by first binning the alignments into 10-kbp non-overlapping windows and calculating the mean sequence identity in each window. Because TandemAligner is only optimized for tandem repeat arrays, we only assessed the sequence identity in the α-satellite HOR array(s) of each centromeric region and did not use it to assess the sequence identity in any other region.

### Pairwise sequence identity heat maps

To generate pairwise sequence identity heat maps of each centromeric region, we ran StainedGlass^55^ (v6.7.0) with the following parameters: window=5000, mm_f=30000, and mm_s=1000. We normalized the color scale across the StainedGlass plots by binning the % sequence identities equally and recoloring the data points according to the binning. To generate heat maps that only show the variation between centromeric regions, we ran StainedGlass^55^ (v6.7.0) with the following parameters: window=5000, mm_f=40000, and mm_s=20000. As above, we normalized the color scale across the StainedGlass plots by binning the % sequence identities equally and recoloring the data points according to the binning.

### Estimation of α-satellite HOR array length

To estimate the length of the α-satellite HOR arrays of each centromere in the CHM1, CHM13, and 56 diverse genome assemblies^7, 18^, we first ran RepeatMasker^54^ (v4.1.0) on the assemblies and identified contigs containing α-satellite repeats, marked by “ALR/Alpha". We extracted these α-satellite-containing contigs and ran Hum-AS-HMMER (https://github.com/fedorrik/HumAS-HMMER_for_AnVIL) on each of them. We subsequently extracted contigs containing “live” α-satellite HORs (denoted with an “L” in the Hum-AS-HMMER BED file). Then, we filtered out contigs that had incomplete α-satellite HOR arrays (e.g., those that did not traverse into unique sequence), thereby limiting our analysis to only complete α-satellite HOR arrays. Additionally, we assessed the integrity of each of the α-satellite HOR array-containing contigs with NucFreq^17^ to ensure that they were completely and accurately assembled, filtering out those with evidence of a deletion, duplication, or misjoin. Finally, we calculated the length of the α-satellite HOR arrays in the remaining contigs by taking the minimum and maximum coordinate of the “live” α-satellite HOR arrays and plotting their lengths with Graphpad Prism (v9).

### Sequence composition and organization of α-satellite HOR arrays

To determine the sequence composition and organization of each α-satellite HOR array in the CHM1, CHM13, and 56 diverse genome assemblies^7, 18^, we ran Hum-AS-HMMER (https://github.com/fedorrik/HumAS-HMMER_for_AnVIL) on centromeric contigs with the default parameters and parsed the resulting BED file with StV (https://github.com/fedorrik/stv). This generated a BED file with each α-satellite HOR sequence composition and its organization along the α-satellite HOR arrays. We used the stv_row.bed file to visualize the organization of the α-satellite HOR arrays with R^56^ (v1.1.383) and the ggplot2 package^57^.

### CpG methylation analysis

To determine the CpG methylation status of each CHM1 centromere, we aligned CHM1 ONT reads >30 kbp in length to the CHM1 whole-genome assembly via Winnowmap^40^ (v1.0) and then assessed the CpG methylation status of the centromeric regions with Nanopolish^58^ (v0.13.3). Nanopolish distinguishes 5-methylcytosines from unmethylated cytosines via a Hidden Markov Model (HMM) on the raw nanopore current signal. The methylation caller generates a log-likelihood value for the ratio of probability of methylated to unmethylated CpGs at a specific *k*-mer. We filtered methylation calls using the nanopore_methylation_utilities tool^59^ (https://github.com/isaclee/nanopore-methylation-utilities), which uses a log-likelihood ratio of 2.5 as a threshold for calling methylation. CpG sites with log-likelihood ratios greater than 2.5 (methylated) or less than −2.5 (unmethylated) are considered high quality and included in the analysis. Reads that do not have any high-quality CpG sites are filtered from the BAM for subsequent methylation analysis. Nanopore_methylation_utilities integrates methylation information into the BAM file for viewing in IGV’s^41^ bisulfite mode, which was used to visualize CpG methylation. To determine the size of hypomethylated region (termed “centromere dip region”, or CDR^26^) in each centromere, we developed a novel tool, CDR-Finder (https://github.com/arozanski97/CDR-Finder). This tool first bins the assembly into 5-kbp windows, computes the median CpG methylation frequency within windows containing α-satellite (as determined by RepeatMasker^54^ (v4.1.0), selects bins that have a lower CpG methylation frequency than the median frequency in the region, merges consecutive bins into a larger bin, filters for merged bins that are >50 kbp, and reports the location of these bins.

### Native CENP-A ChIP-seq and analysis

To determine the location of centromeric chromatin within the CHM1 genome, we performed two independent replicates of native CENP-A ChIP-seq on CHM1 cells as described previously^16^, with some modifications. Briefly, 3-4 x 10^7^ cells were collected and resuspended in 2 mL of ice-cold buffer I (0.32 M sucrose, 15 mM Tris, pH 7.5, 15 mM NaCl, 5 mM MgCl_2_, 0.1 mM EGTA, and 2x Halt Protease Inhibitor Cocktail (Thermo Fisher, 78429)). 2 mL of ice-cold buffer II (0.32 M sucrose, 15 mM Tris, pH 7.5, 15 mM NaCl, 5 mM MgCl_2_, 0.1 mM EGTA, 0.1% IGEPAL, and 2x Halt Protease Inhibitor Cocktail) was added, and samples were placed on ice for 10 min. The resulting 4 mL of nuclei were gently layered on top of 8 mL of ice-cold buffer III (1.2 M sucrose, 60 mM KCl, 15 mM, Tris pH 7.5, 15 mM NaCl, 5 mM MgCl_2_, 0.1 mM EGTA, and 2x Halt Protease Inhibitor Cocktail (Thermo Fisher, 78429)) and centrifuged at 10,000*g* for 20 min at 4°C. Pelleted nuclei were resuspended in buffer A (0.34 M sucrose, 15 mM HEPES, pH 7.4, 15 mM NaCl, 60 mM KCl, 4 mM MgCl_2_, and 2x Halt Protease Inhibitor Cocktail) to 400 ng/mL. Nuclei were frozen on dry ice and stored at 80°C. MNase digestion reactions were carried out on 200-300 μg chromatin, using 0.2-0.3 U/μg MNase (Thermo Fisher, 88216) in buffer A supplemented with 3 mM CaCl_2_ for 10 min at 37°C. The reaction was quenched with 10 mM EGTA on ice and centrifuged at 500*g* for 7 min at 4°C. The chromatin was resuspended in 10 mM EDTA and rotated at 4°C for 2 h. The mixture was adjusted to 500 mM NaCl, rotated for another 45 min at 4°C and then centrifuged at max speed (21,100*g*) for 5 min at 4°C, yielding digested chromatin in the supernatant. Chromatin was diluted to 100 ng/ml with buffer B (20 mM Tris, pH 8.0, 5 mM EDTA, 500 mM NaCl and 0.2% Tween 20) and precleared with 100 μL 50% protein G Sepharose bead (Abcam, ab193259) slurry for 20 min at 4°C, rotating. Precleared supernatant (10-20 μg bulk nucleosomes) was saved for further processing. To the remaining supernatant, 20 μg mouse monoclonal anti-CENP-A antibody (Enzo, ADI-KAM-CC006-E) was added and rotated overnight at 4°C. Immunocomplexes were recovered by the addition of 200 mL 50% protein G Sepharose bead slurry followed by rotation at 4°C for 3 h. The beads were washed 3x with buffer B and once with buffer B without Tween. For the input fraction, an equal volume of input recovery buffer (0.6 M NaCl, 20 mM EDTA, 20 mM Tris, pH 7.5, and 1% SDS) and 1 mL of RNase A (10 mg/mL) was added, followed by incubation for one hour at 37°C. Proteinase K (100 mg/ml, Roche) was then added, and samples were incubated for another 3 h at 37°C. For the ChIP fraction, 300 μL of ChIP recovery buffer (20 mM Tris, pH 7.5, 20 mM EDTA, 0.5% SDS and 500 mg/mL Proteinase K) was added directly to the beads and incubated for 3-4 h at 56°C. The resulting Proteinase K-treated samples were subjected to a phenol-chloroform extraction followed by purification with a Qiagen MinElute PCR purification column. Unamplified bulk nucleosomal and ChIP DNA were analyzed using an Agilent Bioanalyzer instrument and a 2100 High Sensitivity Kit.

Sequencing libraries were generated using the TruSeq ChIP Library Preparation Kit - Set A (Illumina, IP-202-1012) according to the manufacturer’s instructions, with some modifications. Briefly, 5–10 ng bulk nucleosomal or ChIP DNA was end-repaired and A-tailed. Illumina TruSeq adaptors were ligated, libraries were size-selected to exclude polynucleosomes using an E-Gel SizeSelect II agarose gel, and the libraries were PCR-amplified using the PCR polymerase and primer cocktail provided in the kit. The resulting libraries were submitted for 150 bp, paired-end Illumina sequencing using a NextSeq 500/550 High Output Kit v2.5 (300 cycles). The resulting reads were assessed for quality using FastQC (https://github.com/s-andrews/FastQC), trimmed with Sickle (v1.33; https://github.com/najoshi/sickle) to remove low-quality 5’ and 3’ end bases, and trimmed with Cutadapt^60^ (v1.18) to remove adapters.

Processed CENP-A ChIP and bulk nucleosomal reads were aligned to the CHM1 whole-genome assembly using BWA-MEM^61^ (v0.7.17) and the following parameters: bwa mem -k 50 -c 1000000 {index} {read1.fastq.gz} {read2.fastq.gz}. The resulting SAM files were filtered using SAMtools^48^ (v1.9) with flag score 2308 to prevent multi-mapping of reads. With this filter, reads mapping to more than one location are randomly assigned a single mapping location, thereby preventing mapping biases in highly identical regions. Alignments were normalized with deepTools^62^ (v3.4.3) bamCompare with the following parameters: bamCompare -b1 {ChIP.bam} -b2 {bulk_nucleosomal.bam} --operation ratio -- binSize 1000 -o {out.bw}.

### Human and NHP α-satellite suprachromosomal family (SF) classification and strand analysis

To determine the α-satellite SF content and strand orientation of human and NHP centromeres, we generated custom SF and strand tracks for each centromere assembly in the UCSC Human Genome Browser. For the CHM1 centromeres, we built two additional tracks: one showing each α-satellite monomer belonging to known human HORs (ASat-HOR track) and another showing structural variation in human HORs (StV track). All tracks were built and color-coded as described previously^4^ and are publicly available at the following URLs: https://genome.ucsc.edu/s/fedorrik/chm1_cen (CHM1); https://genome.ucsc.edu/s/fedorrik/T2T_dev (CHM13); https://genome.ucsc.edu/s/fedorrik/cen_primates (chimpanzee, orangutan, and macaque). We note that the SF annotation coverage in macaque is sometimes discontinuous (some monomers are not annotated due to significant divergence of macaque dimers from their progenitor Ka class monomers). However, most monomers are identified as Ka, which indicates SF7. In orangutan centromeres, most monomers are identified as R1 and R2, which indicates SF5. In chimpanzee and human autosome and X chromosome centromeres, active arrays are formed by J1 and J2 (SF1), D1, FD and D2 (SF2), and W1-W5 (SF3) monomers. The only exception uncovered in this paper is the centromere of chimpanzee chromosome 5, which appears to be formed by R1 and R2 (SF5), with some monomers identified as J4 and Ga. The former belongs to SF01, which represents the generation of α-satellite intermediate between the progenitor SF5 and the more derived SF1, and J4 is particularly close to the R1 monomer. Also, the other SF01 monomers, such as J3, J5 and J6, are absent in the array, which indicates that it is not genuine SF01. Therefore, the J4 monomer in chimpanzee centromere 5 should be considered variant R1. Similarly, occasional Ga monomers belong to SF4, which is the direct progenitor of SF5, and Ga is very close to R2. Therefore, Ga monomers dispersed in the SF5 array are just misclassed R2 monomers. Thus, the whole chimpanzee chromosome 5 α-satellite HOR array should be classified as SF5, despite the abovementioned contaminations.

### Human and NHP phylogenetic analysis

To assess the phylogenetic relationship between α-satellite repeats in human and NHP genomes, we first masked every non-α-satellite repeat in the CHM1, CHM13, HG00733, chimpanzee, orangutan, and macaque centromere assemblies using RepeatMasker^54^ (v4.1.0). Then, we subjected the masked assemblies to StringDecomposer^63^ using α-satellite monomers derived from the T2T-CHM13 reference genome^3^ (v2.0). This tool identifies the location of α-satellite monomers in the assemblies, and we used this to extract the α-satellite monomers from the HOR/dimeric array and monomeric regions into multi-FASTA files. We randomly selected 100 and 50 α-satellite monomers from the HOR/dimeric array and monomeric regions, respectively, and aligned them with MAFFT^64, 65^ (v7.453). We used IQ-TREE^66^ (v2.1.2) to reconstruct the maximum-likelihood phylogeny with model selection and 1000 bootstraps. The resulting tree file was visualized in iTOL^67^

To estimate sequence divergence along the pericentromeric regions, we first mapped each NHP centromere assembly to the CHM13 centromere assembly using minimap2^39^ (v2.17-r941) with the following parameters: -ax asm20 --eqx -Y -t 8 -r 500000. Then, we generated a BED file of 10 kbp windows located within the CHM13 centromere assembly. We used the BED file to subset the BAM file, which was subsequently converted into a set of FASTA files. FASTA files contained at least 5 kbp of orthologous sequences from one or more NHP centromere assemblies. Pairs of human and NHP orthologous sequences were realigned using MAFFT^64, 65^ (v7.453) and the following command: mafft -- maxiterate 1000 --localpair. Sequence divergence was estimated using the Tamura-Nei substitution model^68^, which accounts for recurrent mutations and differences between transversions and transitions as well as within transitions. Mutation rate per segment was estimated using Kimura’s model of neutral evolution^69^. In brief, we modeled the estimated divergence (D) is a result of between-species substitutions and within-species polymorphisms; i.e.,

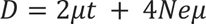

where Ne is the ancestral human effective population size, t is the divergence time for a given human– NHP pair, and μ is the mutation rate. We assumed a generation time of [20, 29] years and the following divergence times: human–macaque = [23e6, 25e6] years, human–orangutan = [12e6, 14e6] years, human–chimpanzee = [4e6, 6e6] years. To convert the genetic unit to a physical unit, our computation also assumes Ne=10,000 and uniformly drawn values for the generation and divergence times.

### Human-specific phylogenetic analysis

To determine the phylogenetic relationship and divergence times between centromeric regions from chromosomes 5, 7, and 10-14 in the CHM1, CHM13, and 56 other diverse human genomes (sequenced and assembled by the HPRC^7^ and HGSVC^18^), we first identified contigs with complete and accurately assembled centromeric α-satellite HOR arrays, as determined by RepeatMasker^54^ (v4.1.0) and NucFreq^17^ analysis. Then, we aligned each of these contigs to the T2T-CHM13 reference genome^3^ (v2.0) via minimap2^39^ (v2.24). We also aligned the chimpanzee whole-genome assembly to the T2T-CHM13 reference genome^3^ (v2.0) to serve as an outgroup in our analysis. We identified 20-kbp regions in the flanking monomeric α-satellite or unique regions on the p- or q-arms and ensured that the region we had selected had only a single alignment from each haplotype to the reference genome. Then, we aligned these regions to each other using MAFFT^64, 65^ (v7.453) and the following command: mafft –auto –thread {num_of_threads} {multi-fasta.fasta}. We used IQ-TREE^66^ (v2.1.2) to reconstruct the maximum-likelihood phylogeny with model selection and 1000 bootstraps. The resulting tree file was visualized in iTOL^67^. Clusters of α-satellite HOR arrays with a single monophyletic origin were assessed for gains and loss of α-satellite base pairs, monomers, HORs, and distinct structural changes manually.

## DATA AVAILABILITY

All datasets generated and/or used in this study are publicly available and listed in **Extended Data Table 10** with their BioProject ID, accession # (if available), and/or URL. For convenience, we also list the BioProject IDs and/or URLs here: CHM1 whole-genome assembly with complete centromeres (PRJNA975207); CHM1 PacBio HiFi data (PRJNA726974); CHM1 ONT data (PRJNA869061); CHM Illumina data (PRJNA246220); CHM1 Strand-Seq alignments (https://doi.org/10.5281/zenodo.7959305); CHM1 CENP-A ChIP-seq data (PRJNA975217); HG00733 PacBio HiFi data (PRJNA975575 and PRJEB36100); HG00733 ONT data (PRJNA975575, PRJNA686388, and PRJEB37264); and NHP [chimpanzee (Clint; S006007), orangutan (Susie; PR01109), and macaque (AG07107)] PacBio HiFi and ONT data (PRJNA659034).

## CODE AVAILABILITY

Custom code for the sequence assembly of primate centromeric regions is available at https://github.com/arozanski97/SUNK-based-contig-scaffolding. Custom code to detect hypomethylated regions within centromeric regions, termed centromere dip regions” or CDRs^26^, is available at https://github.com/arozanski97/CDR-Finder. All other code is publicly available.

## ACKNOWLEDGMENTS

We thank Google, Inc. for basecalling CHM1 PacBio circular consensus sequencing (CCS) data with DeepConsensus; P.M. Lansdorp (BC Cancer Institute and UBC) and D.C.J. Spierings (European Research Institute for the Biology of Aging) for generating and sharing the CHM1 Strand-seq data; P. Hsieh (UMN) for assistance with phylogenetic analyses; Viviane Slon (Tel-Aviv University) for providing space and resources for the α-satellite suprachromosomal family and strand analysis with funding from the Genetic Society of America and the Gruber foundation; Z. Zhao for comments and suggestions on figures; and T. Brown (UW) for assistance in editing this manuscript. This research was supported, in part, by funding from the National Institutes of Health (NIH) National Human Genome Research Institute (NHGRI) R01 HG010169 (EEE); National Institute of General Medical Sciences (NIGMS) K99 GM147352 (GAL); National Cancer Institute (NCI) R01 CA266339 (JLG); Intramural Research Program of the National Human Genome Research Institute (NHGRI) at NIH (MR, SK, SN, AMP); Shanghai Jiao Tong University 2030 Program WH510363001-7 (YM); and Center for Integration in Science of the Ministry of Aliyah, Israel (IAA). This work utilized the computational resources of the NIH HPC Biowulf cluster (https://hpc.nih.gov). EEE is an investigator of the Howard Hughes Medical Institute.

## AUTHOR CONTRIBUTIONS

GAL and EEE conceived the project; GAL, JKL, KH, and KMM generated sequencing data; GAL and ANR analyzed sequencing data and performed QC analyses; GAL and ANR generated and validated the CHM1 centromere assemblies; MR, SK, SN, and AMP generated the Verkko CHM1 centromere assemblies; GAL, ANR, FR, YS, and IAA analyzed the CHM1 centromere assemblies; TP performed spectral karyotyping and fluorescent *in situ* hybridization experiments; DP performed Strand-seq analyses; GAL, ANR, and YM performed phylogenetic analyses; JGL, AMP, IAA, and EEE supervised experiments and analyses; GAL, DP, IAA, and EEE developed figures; and GAL, IAA, and EEE drafted the manuscript. All authors have read and approved the manuscript.

