## Extended Data for "The variation and evolution of complete human centromeres"

**This PDF file includes:**

1. **Extended Data Figures 1 to 20**
2. **Extended Data Tables 1 to 10**
3. **References**

### EXTENDED DATA FIGURES

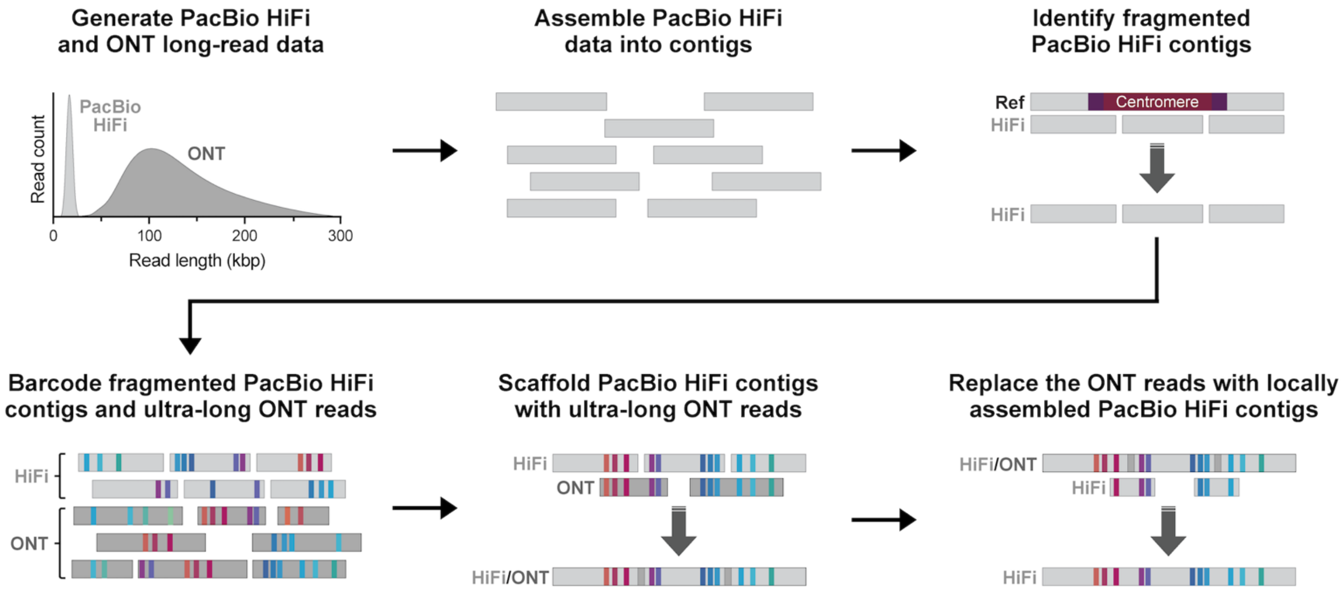


**Extended Data Figure 1. Centromere assembly method.** To assemble each CHM1 centromere, we first generated ~56-fold sequence coverage of Pacific Biosciences (PacBio) high-fidelity (HiFi) data and ~100-fold sequence coverage of Oxford Nanopore Technologies (ONT) data from the CHM1 genome. Then, we assembled the PacBio HiFi data into contigs using an established assembler, hifiasm[1]. We identified PacBio HiFi contigs that were fragmented over the centromeres by aligning them to the T2T-CHM13 reference genome. We barcoded the fragmented centromeric PacBio HiFi contigs and ultra-long (>100 kbp) ONT reads with singly unique nucleotide *k*-mers (SUNKs), creating unique SUNK barcodes. We ordered, oriented, and joined the PacBio HiFi contigs together with ultra-long ONT reads based on shared SUNK barcodes, generating a hybrid PacBio HiFi/ONT-based sequence assembly of each centromere. To improve the base accuracy of each assembly, we replaced the ONT reads with PacBio HiFi contigs that had been locally assembled with HiCanu[2], generating gapless sequence assemblies of each CHM1 centromere that are estimated to be >99.9999% accurate (as determined with Merqury[3]).

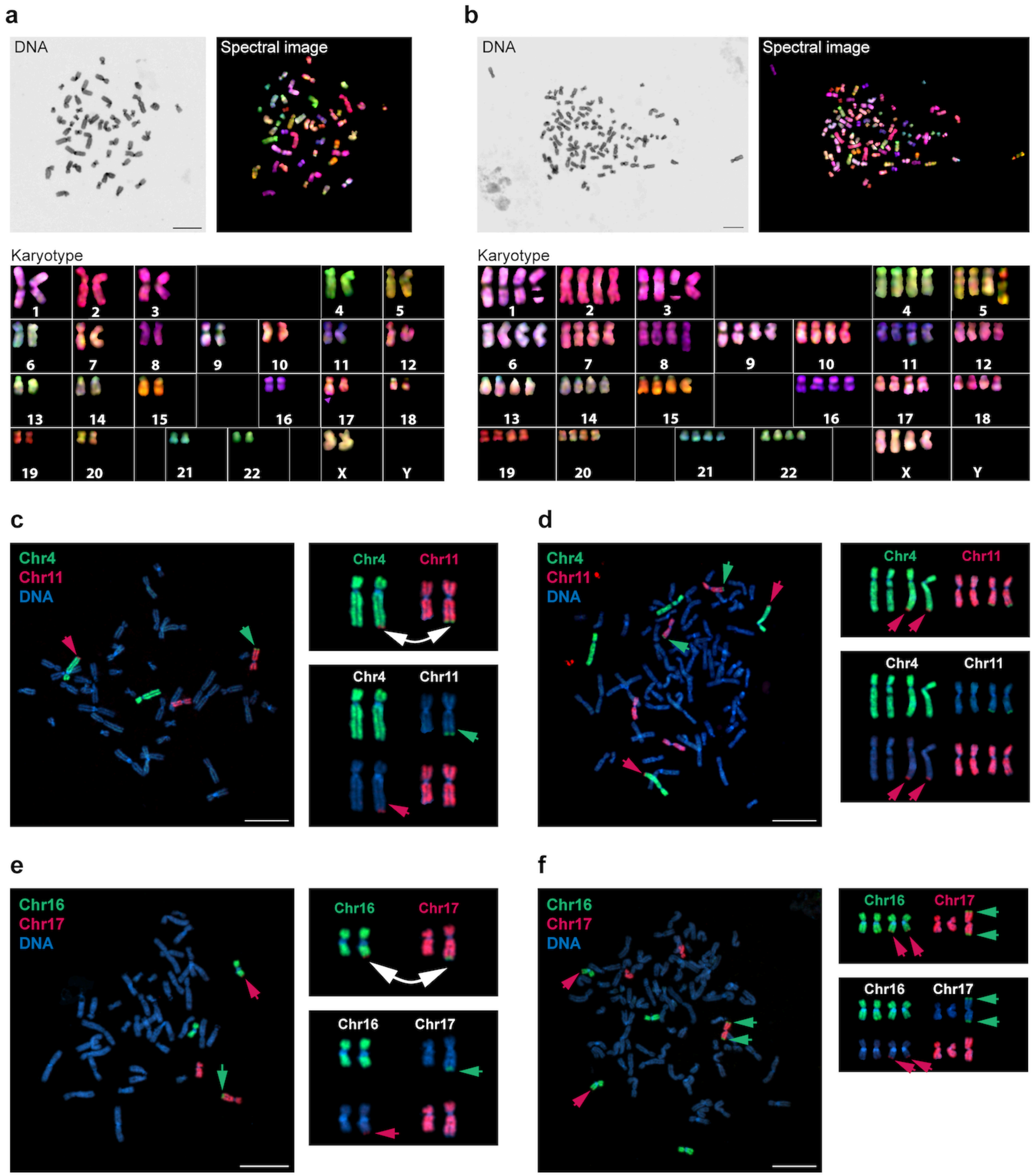


**Extended Data Figure 2. Karyotype of the CHM1 genome. a,b)** Giemsa staining and spectral karyotyping of the CHM1 cell line reveals that approximately 71% of CHM1 cells are in **a)** a diploid/near-diploid state, and 29% of cells are in **b)** a tetraploid/near-tetraploid state. **c-f)** Fluorescent *in situ* hybridization on CHM1 metaphase chromosome spreads reveals that almost all cells have a reciprocal translocation between **c,d)** chromosomes 4q35.1 and 11q24.3 and **e,f)** chromosomes 16q23.3 to 17q25.3. Additionally, 44% of diploid/near-diploid cells and 83% of tetraploid/near-tetraploid cells are missing one copy of chromosome 17 or the chromosome 17 p-arm. Bar, 10μm.

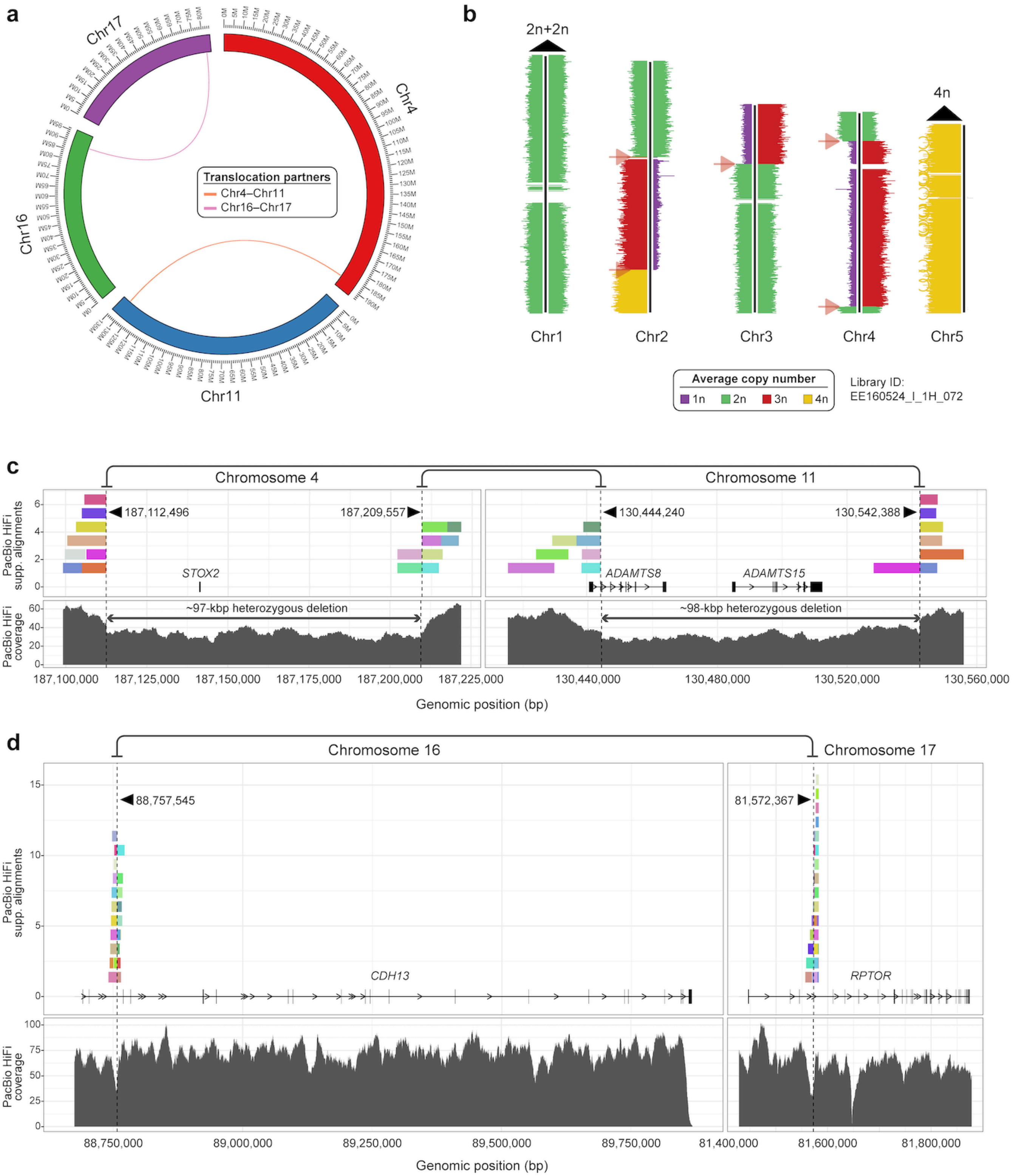


**Extended Data Figure 3. Translocations in the CHM1 genome. a)** Single-cell sequencing of template DNA strands (Strand-seq) from the CHM1 genome confirms the presence of two reciprocal translocations between chromosomes 4q35.1/11q24.3 and 16q23.3/17q25.3 and further refines the breakpoints to chr4:187112496/chr11:130542388, chr4:187209555/chr11:130444240, and chr16:88757545/chr17:81572367 (in T2T-CHM13 v2.0). We note that there are two breakpoints for the chromosome 4q35.1/11q24.3 reciprocal translocation because it is accompanied by a ~97-98 kbp deletion at chr4:187112495-187209555 and chr11:130444240-130542388 (in T2T-CHM13 v2.0). **b)** Example of a CHM1 Strand-seq library mapped to a subset of chromosomes in the T2T-CHM13 reference genome[4]. Each chromosome is depicted as a vertical ideogram, and the distribution of directional sequencing reads is represented by horizontal lines along each chromosome. Reads mapped to the plus strand of the reference genome are shown on the left, and those mapped to the minus strand on the right of each ideogram. The average copy number of each chromosomal region is indicated. On chromosomes 1 and 5, for example, we find an average of 4n copies. On chromosomes 2, 3, and 4, we find low-frequency switches in strand-state (so-called sister-chromatid exchange events, or SCEs[5]) marked by arrows. Nevertheless, the overall copy number across each chromosome sums to 4n. **c, d)** Mapping of CHM1 PacBio HiFi reads to the T2T-CHM13 reference genome[4] reveals the precise breakpoints of the reciprocal translocation between chromosomes **c)** 4q35.1/11q24.3 and **d)** 16q23.3/17q25.3. CHM1 PacBio HiFi reads spanning the translocations are uniquely colored, and predicted translocation breakpoints are indicated with vertical dashed lines. The exon structure of all genes and the read depth of the CHM1 PacBio HiFi data are shown. The chromosome 4q35.1/11q24.3 translocation is associated with a deletion in both chromosomes, resulting in deletion of the *STOX2* and *ADAMTS15* genes and partial deletion of the *ADAMTS8* gene. The chromosome 16q23.3/17q25.3 is associated with a novel fusion of the *CDH13* and *RPTOR* genes.

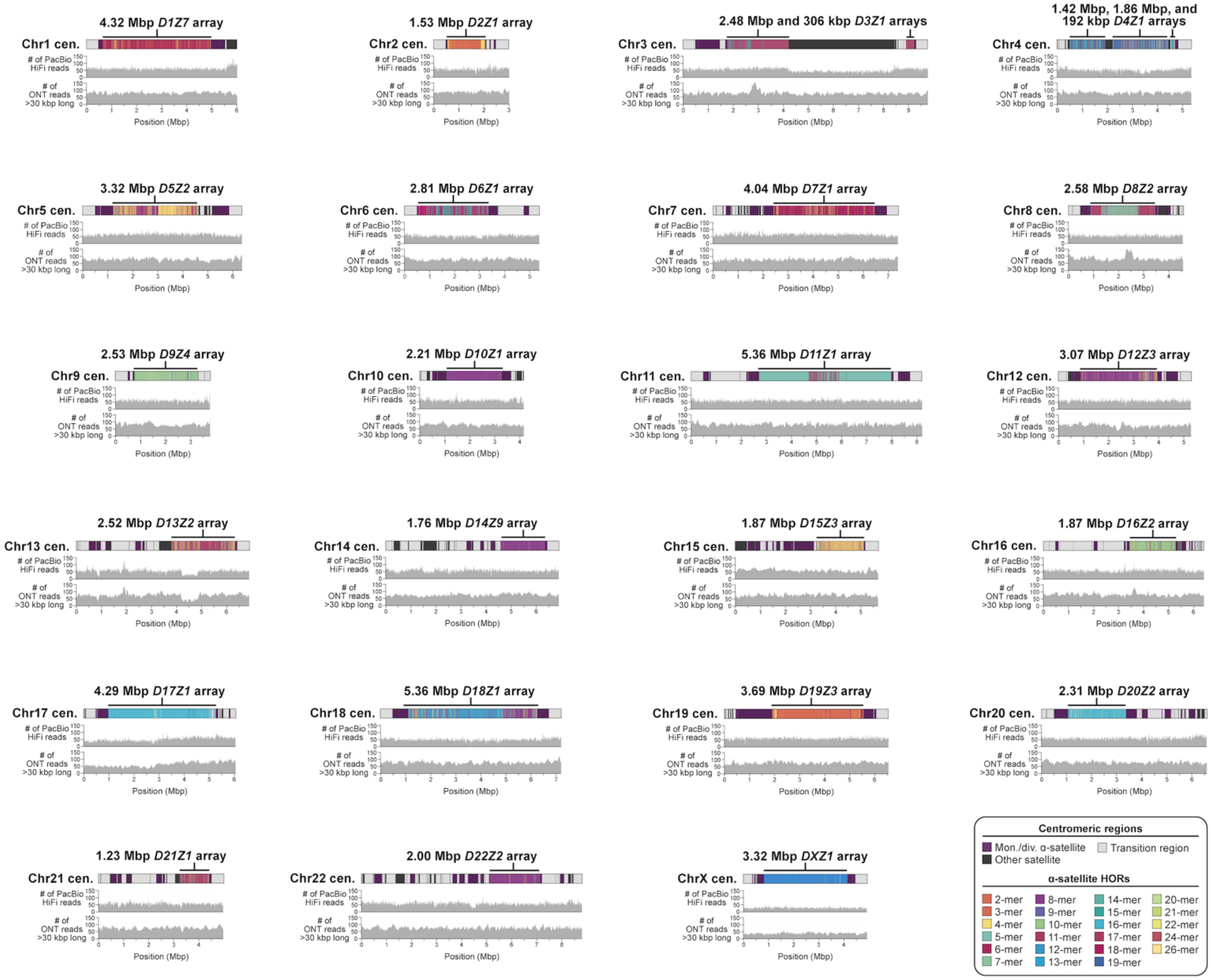


**Extended Data Figure 4. Read-depth profiles of each CHM1 centromere.** Alignment of CHM1 PacBio HiFi and ONT long-read sequencing data to the CHM1 centromere assemblies shows uniform read depth, indicating a lack of large structural errors. However, we note two exceptions that have a biological basis: first, the chromosome 17 centromere has reduced coverage near its p-arm, consistent with the loss of the p-arm in a subset of cells; second, the chromosome 13 centromere has reduced coverage over a 631-kbp region in the core of the centromere, suggesting a deletion of this region in a subset of cells. Both events are supported by CHM1 PacBio HiFi and ONT data.

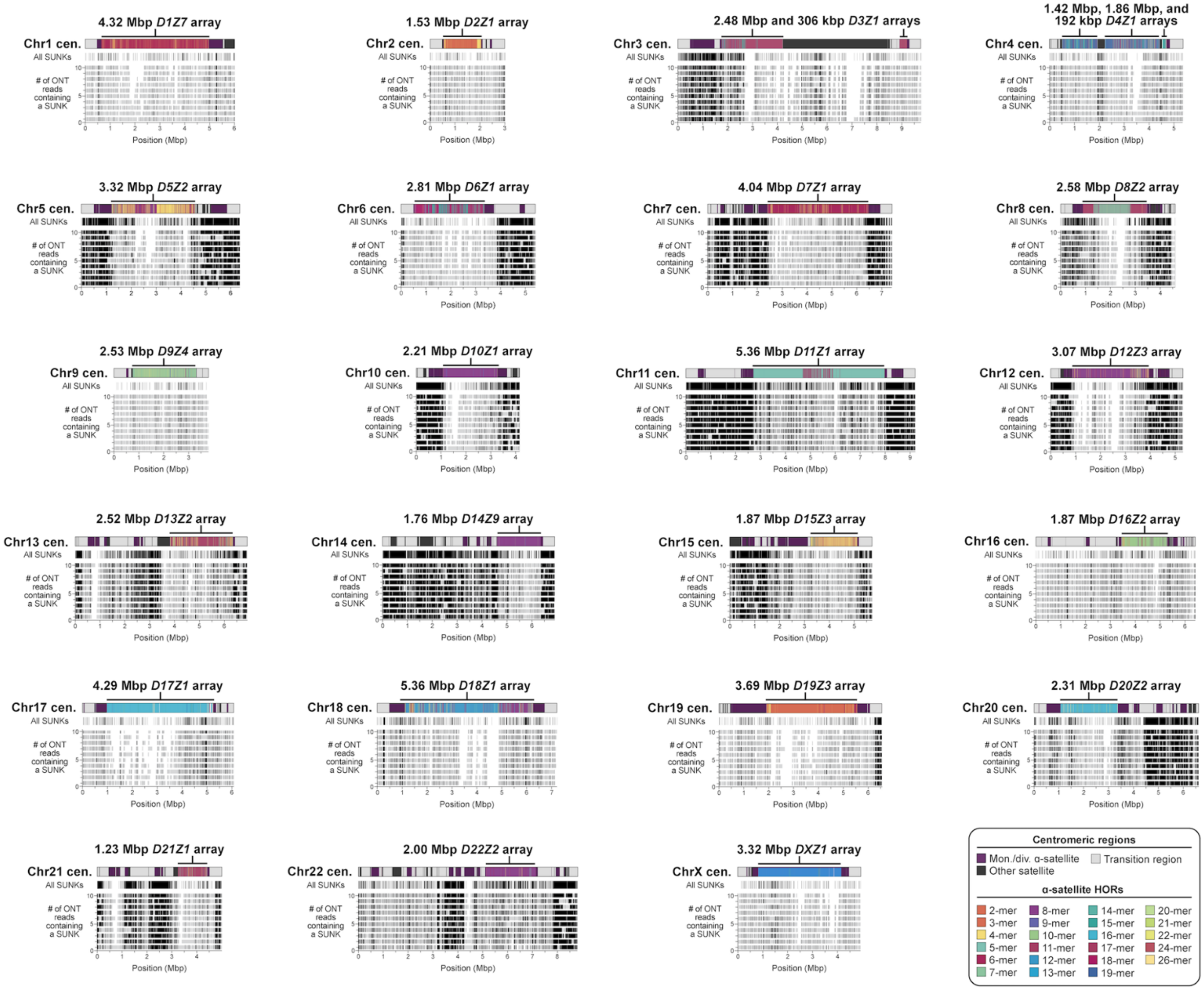


**Extended Data Figure 5. Validation of each CHM1 centromere assembly with native ONT reads via GAVISUNK.** Plots showing the concordance between the CHM1 centromere assemblies and native ONT reads based on patterns of SUNKs (black vertical bars). Plots are generated with GAVISUNK[6].

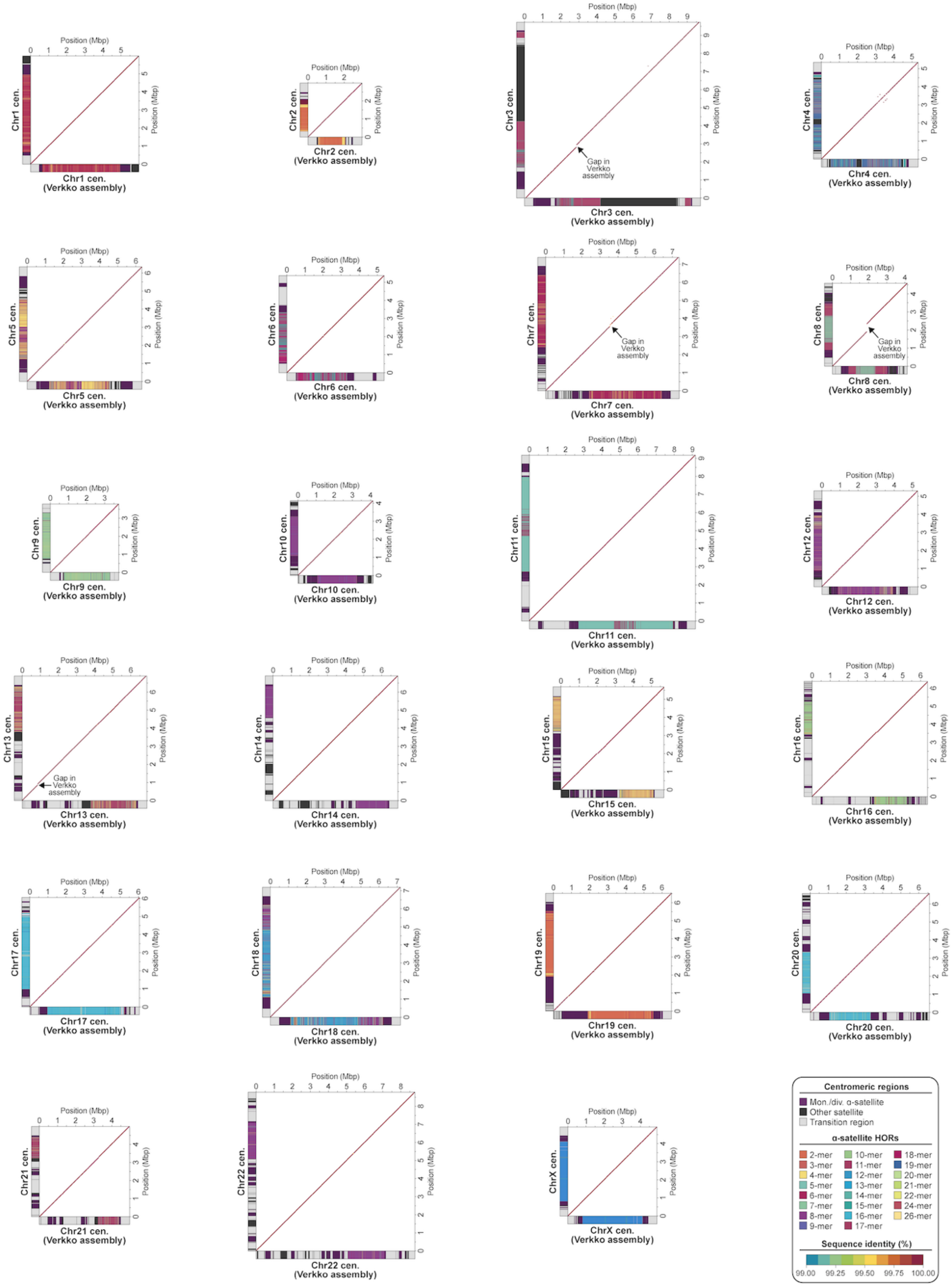


**Extended Data Figure 6. Comparison of each CHM1 centromere assembly to those generated by another assembler, Verkko.** Plots showing the % sequence identity between the CHM1 centromere assemblies generated in this study and those generated via Verkko[7]. Each centromere is >99.9% identical in sequence. Gaps in the Verkko centromere assemblies are indicated. Plots were generated with StainedGlass[8].

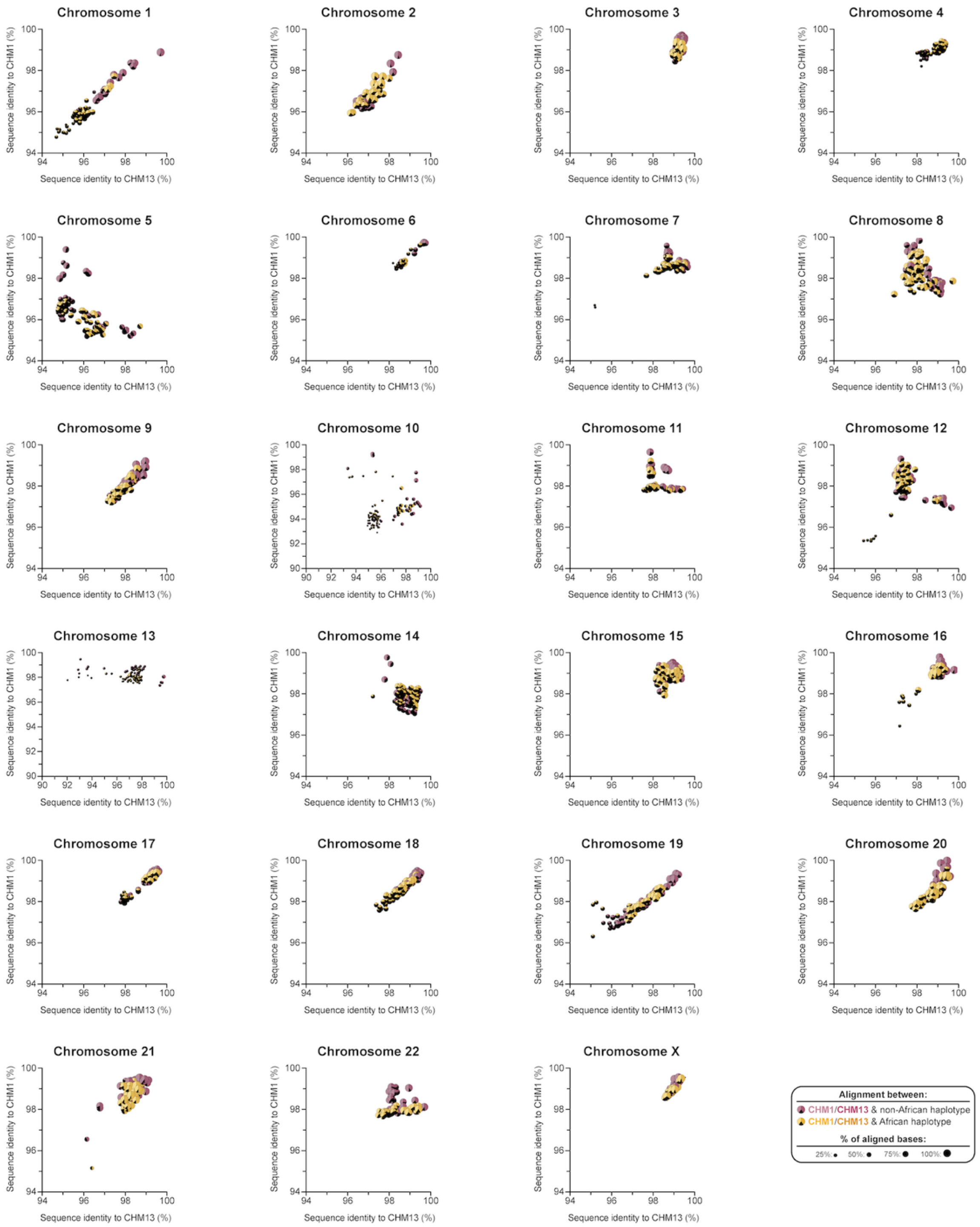


**Extended Data Figure 7. Variation in the sequence and structure of centromeric α-satellite higher-order repeat (HOR) arrays among 56 diverse human genomes.** Plots showing the percent sequence identity between centromeric α-satellite HOR arrays from CHM1 (y-axis), CHM13 (x-axis), and 56 other diverse human genomes [generated by the Human Pangenome Reference Consortium (HPRC)[9] and Human Genome Structural Variation Consortium (HGSVC)[10]]. Each data point shows the percent of aligned bases from each human haplotype to either the CHM1 (left) or CHM13 (right) α-satellite HOR array(s). The percent of unaligned bases are shown in black. The size of each data point corresponds to the total percent of aligned bases among the CHM1 and CHM13 centromeric α-satellite HOR arrays.

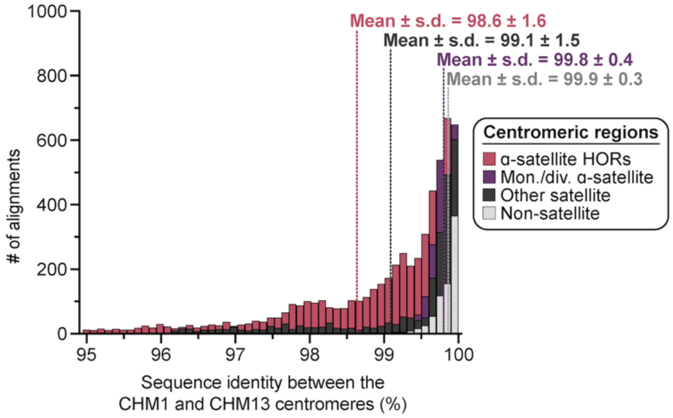


**Extended Data Figure 8. Sequence identities between the CHM1 and CHM13 centromeric regions.** Histogram showing the distribution of sequence identities from complete contig alignments between centromeric regions in the CHM1 and CHM13 genomes. The α-satellite HOR, monomeric/divergent α-satellite, other satellite, and non-satellite portions were assessed separately and reveal a much larger distribution in sequence identities for the α-satellite HORs. The mean and standard deviation (s.d.) are indicated.

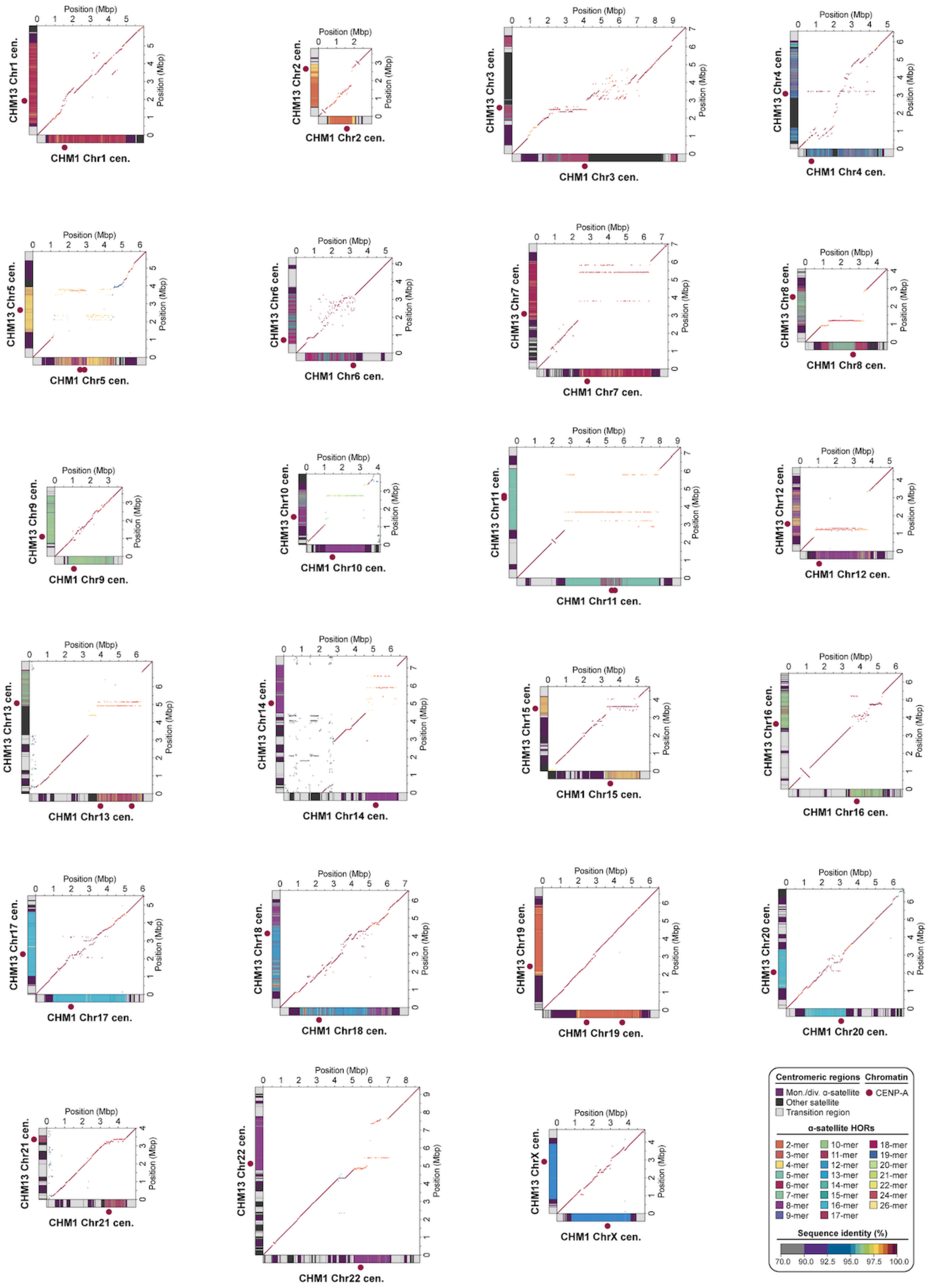


**Extended Data Figure 9. Comparison of the CHM1 and CHM13 centromeric regions.** Dot plots showing the percent sequence identity between the CHM1 and CHM13 centromeric regions. Plots were generated with StainedGlass[8].

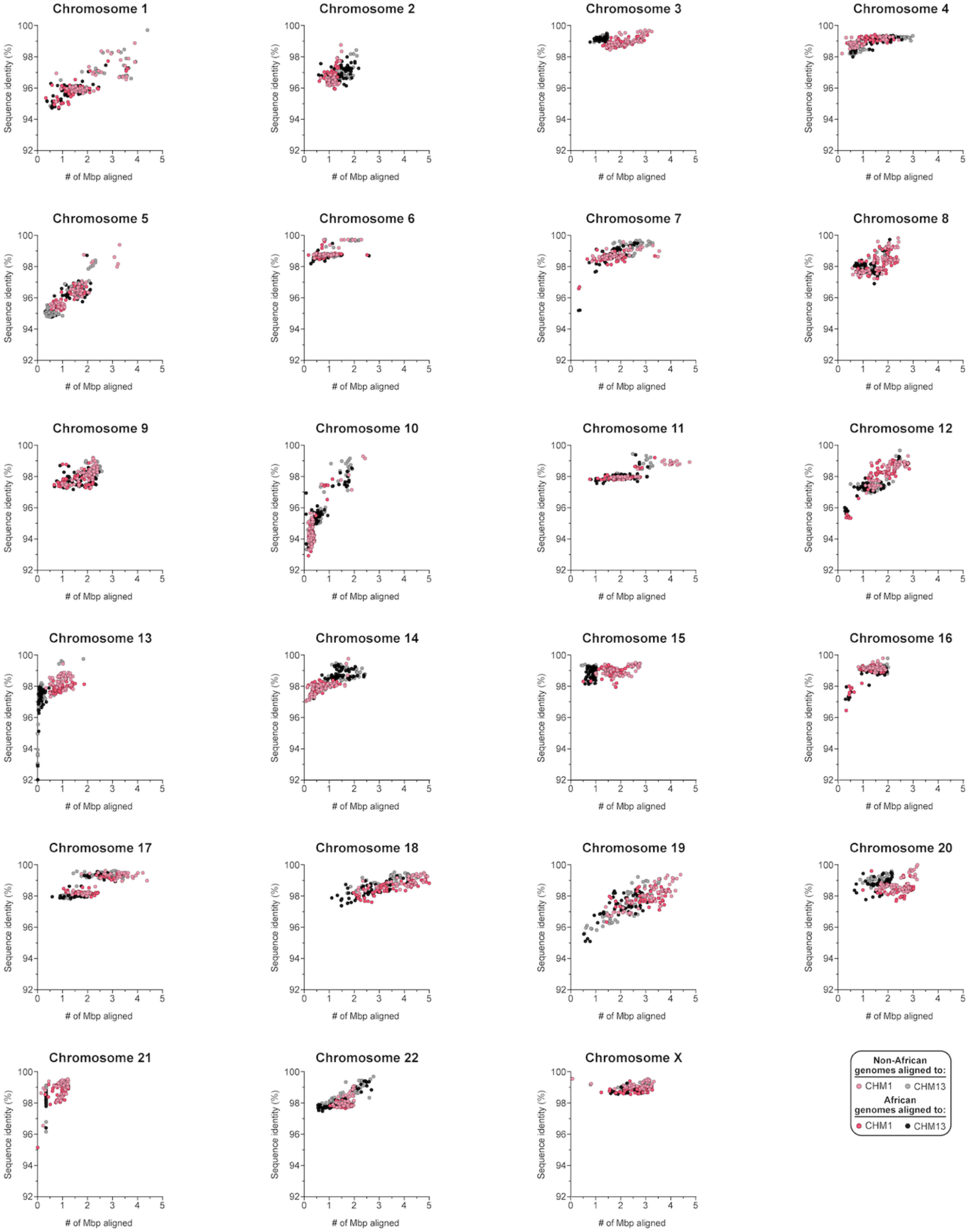


**Extended Data Figure 10. Comparison of CHM1 and CHM13 centromeric α-satellite HOR arrays to those from 56 diverse human genomes.** Plots showing the percent sequence identity and number of megabase pairs (Mbp) aligned for 56 diverse human genomes (112 haplotypes), generated by the HPRC[9] and HGSVC[10], mapped to the CHM1 and CHM13 centromeric regions. Note that each data point represents a haplotype with 1:1 best mapping, although many of the centromeres are not yet complete in the HPRC and HGSVC assemblies.

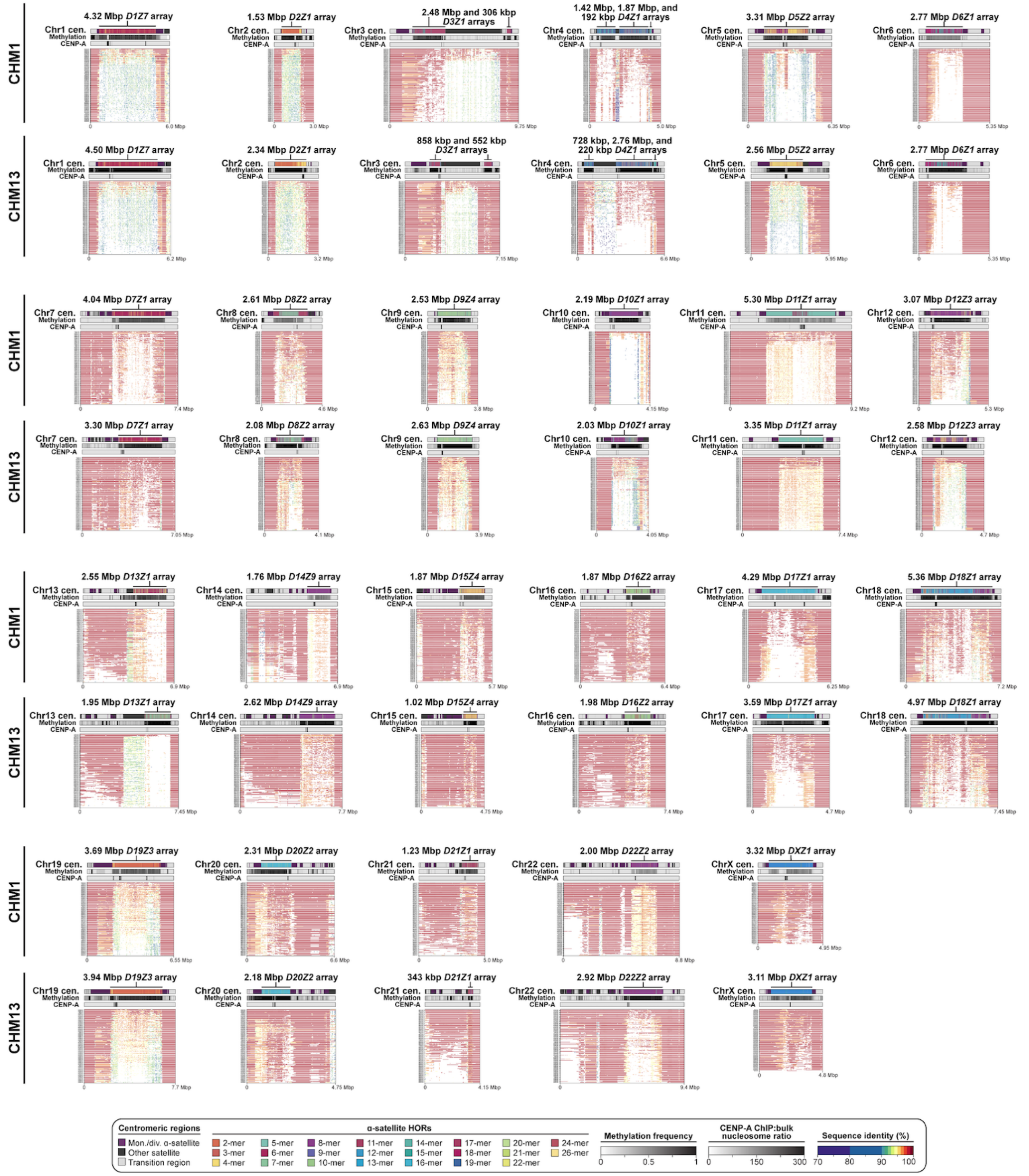


**Extended Data Figure 11. Comparison of the sequence and structure of centromeric regions from the CHM1, CHM13, and 56 diverse human genomes.** Plots showing the sequence organization (top track), CpG methylation frequency (second track), CENP-A nucleosome enrichment (third track), and percent sequence identity of contigs from 56 diverse human genomes (112 haplotypes[9,10]) relative to the CHM1 and CHM13 genomes. Approximately half of all centromeric regions have conserved sequences coinciding with the site of DNA hypomethylation and CENP-A nucleosome enrichment.

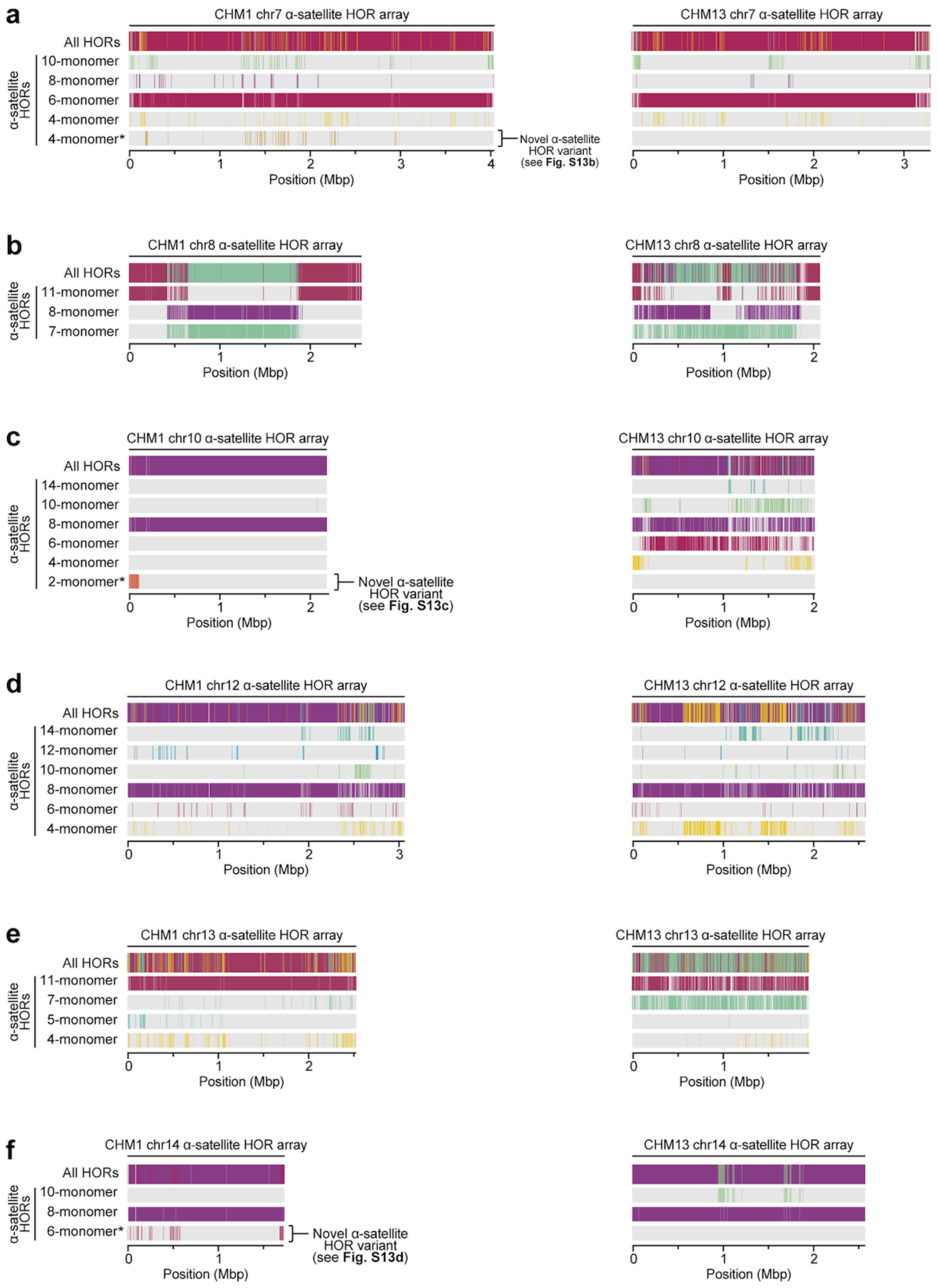


**Extended Data Figure 12. Variation in the sequence and structure of the α-satellite HOR arrays between the CHM1 and CHM13 centromeres. a-h)** Structure of the CHM1 (left) and CHM13 (right) α-satellite HOR arrays from chromosomes **a)** 7, **b)** 8, **c)** 10, **d)** 12, **e)** 13, and **f)** 14. Novel α-satellite HOR variants are indicated.

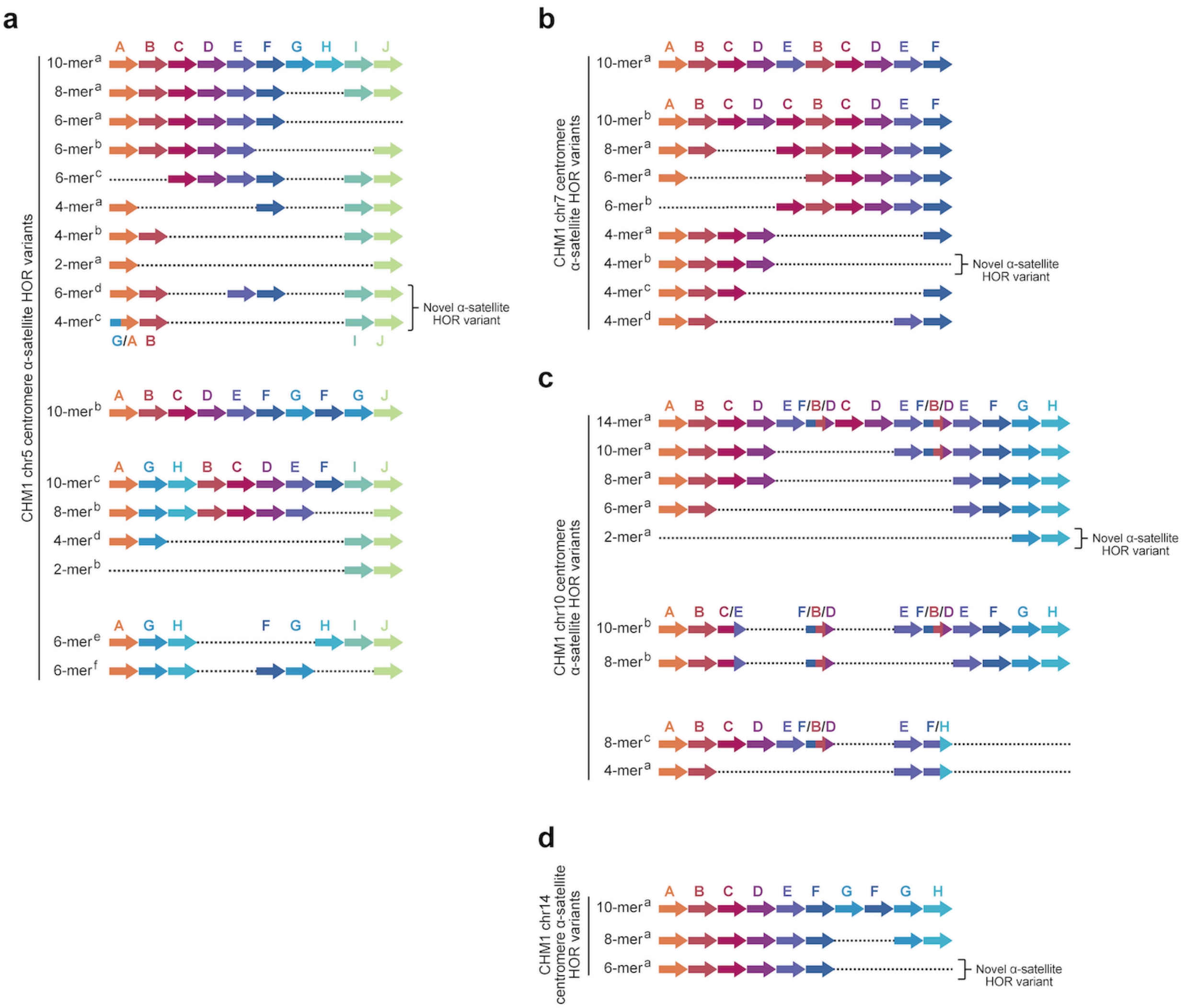


**Extended Data Figure 13. Novel α-satellite HOR variants within the CHM1 centromeres. a-d)** Structures of the α-satellite HOR variants within the CHM1 centromeres from chromosomes **a)** 5, **b)** 7, **c)** 10, and **d)** 14. Novel α-satellite HOR variants are indicated.

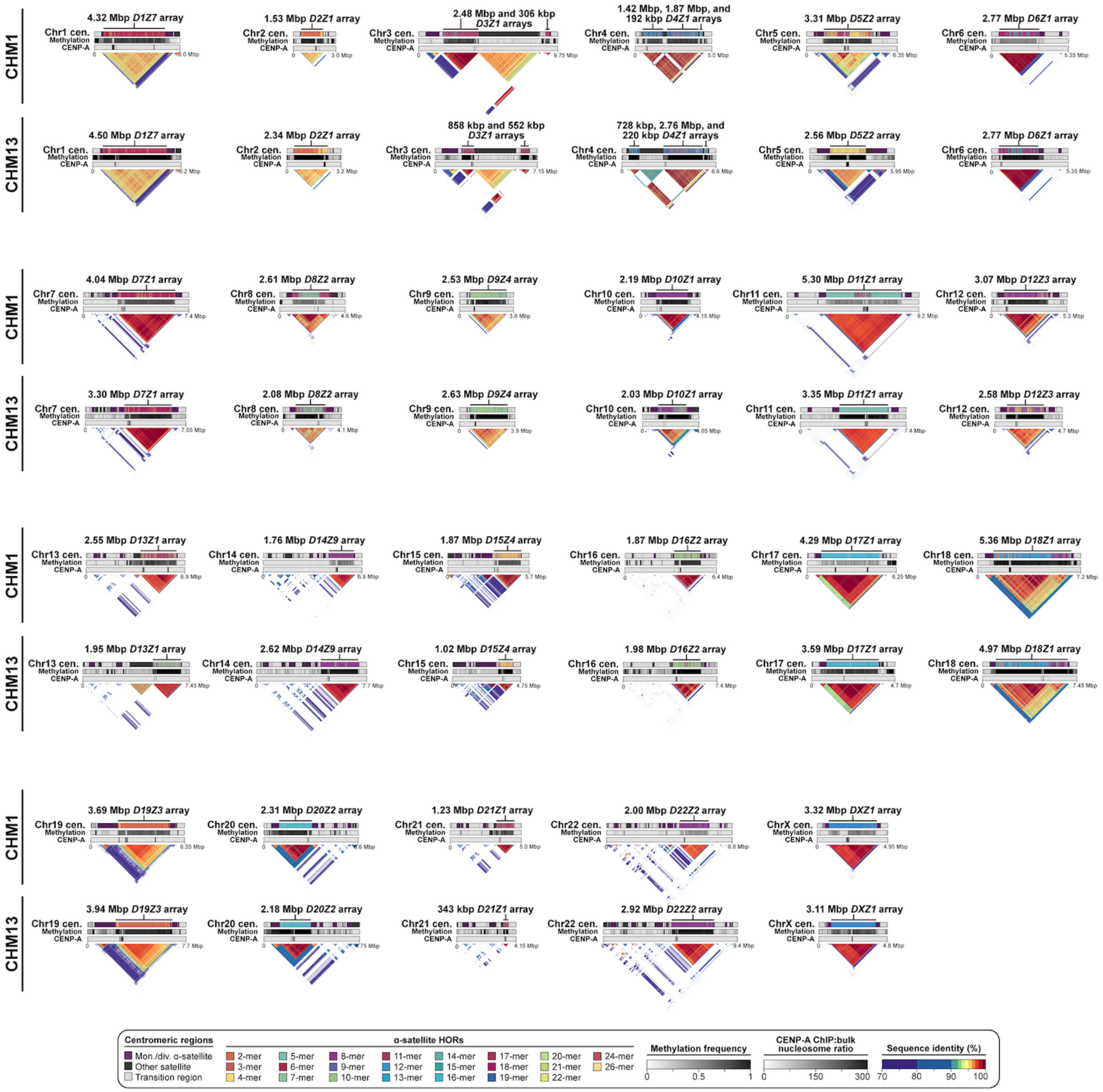

**Extended Data Figure 14. Comparison of the genetic, epigenetic, and evolutionary landscapes between the CHM1 and CHM13 centromeric regions.** Plots showing the sequence organization (top track), CpG methylation frequency (second track), CENP-A nucleosome enrichment (third track), and evolutionary layers (bottom triangle) for each CHM1 and CHM13 centromeric region.

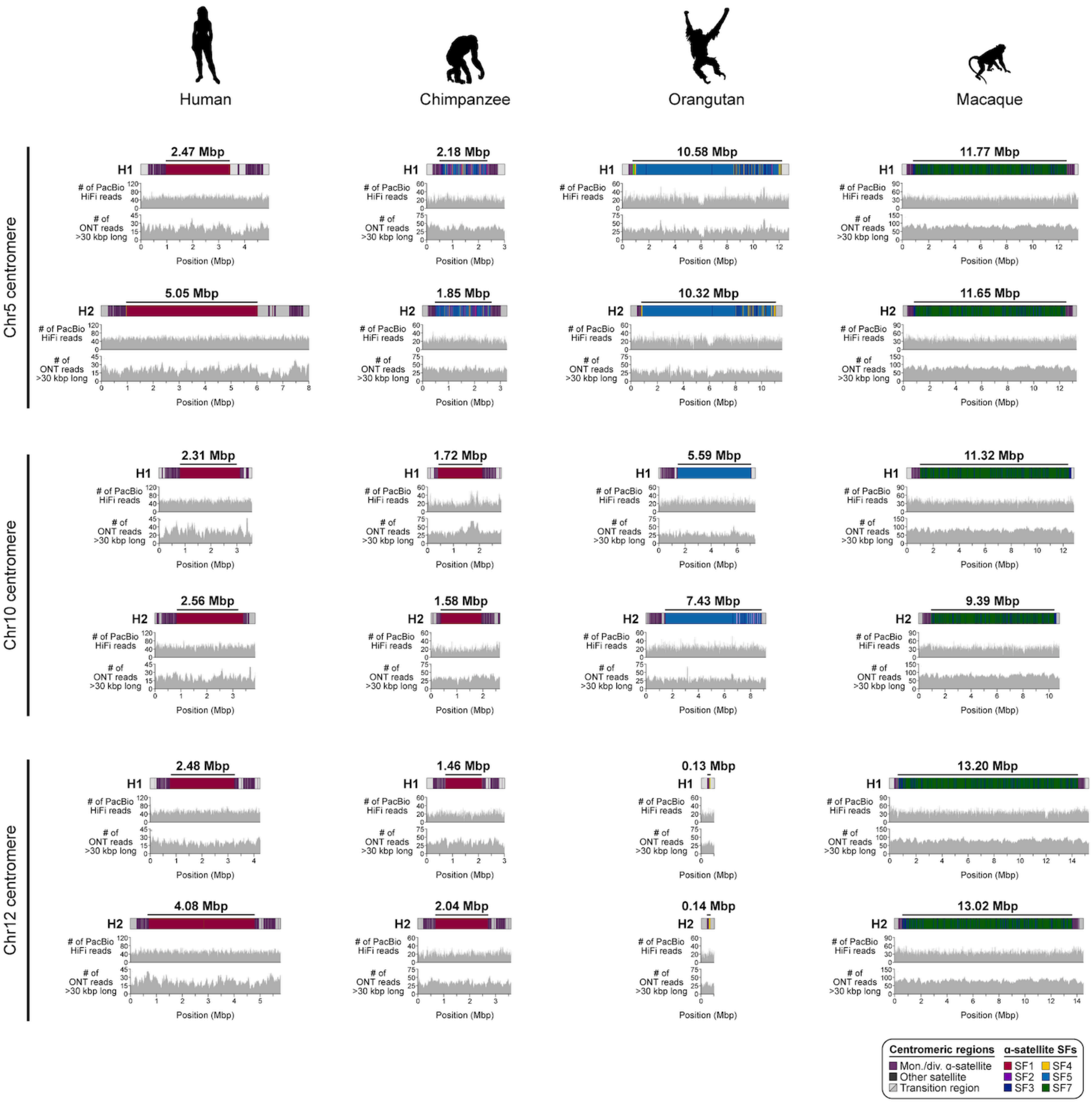


**Extended Data Figure 15. Read-depth profiles of the chromosome 5, 10, and 12 centromeric regions from the human, chimpanzee, orangutan, and macaque genomes.** Alignment of PacBio HiFi and ONT long-read sequencing data to the centromere assemblies from diverse primate species shows uniform read depth, indicating a lack of large structural errors. The human genome is HG00733.

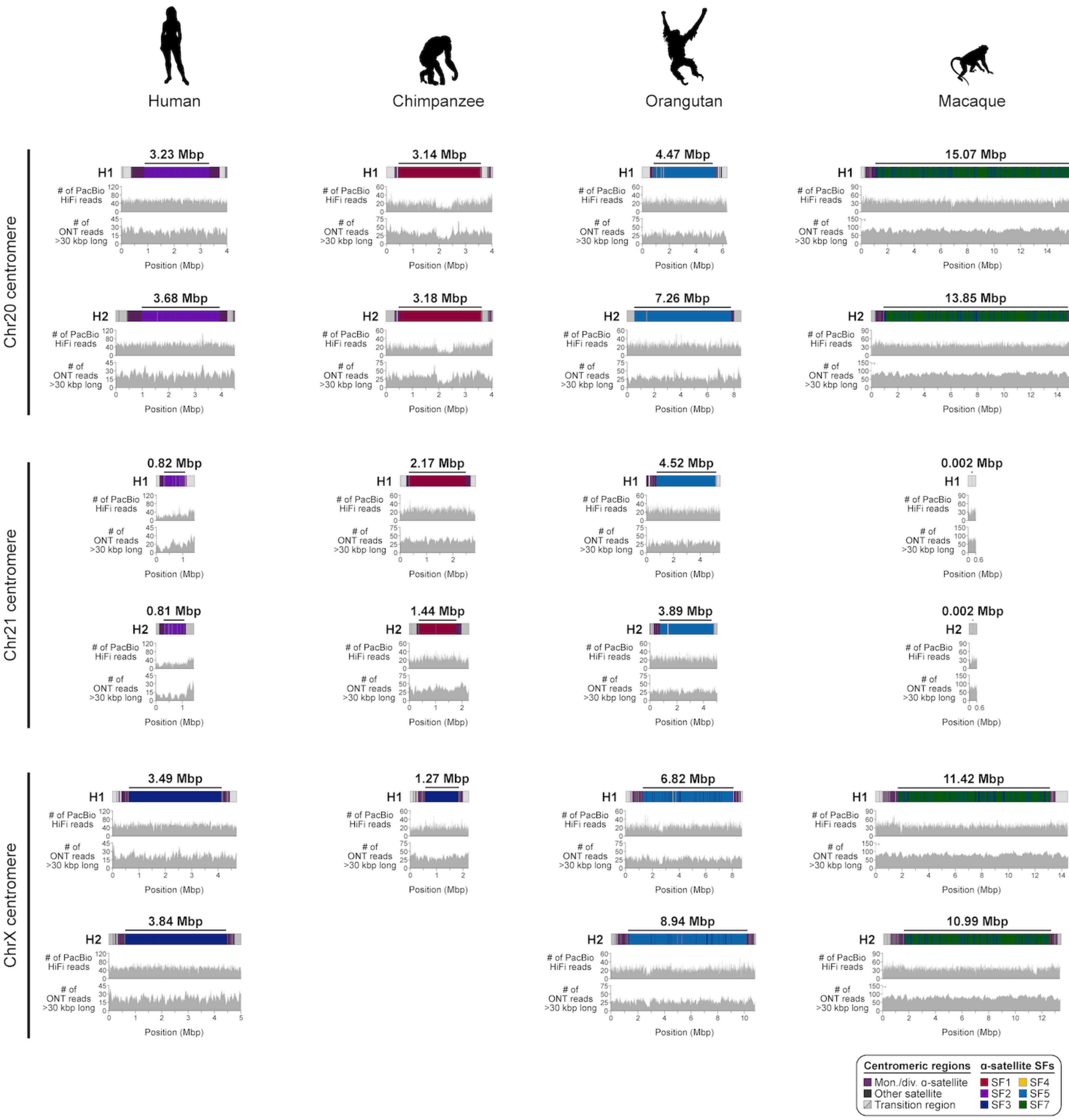


**Extended Data Figure 16. Read-depth profiles of the chromosome 20, 21, and X centromeric regions from the human, chimpanzee, orangutan, and macaque genomes.** Alignment of PacBio HiFi and ONT long-read sequencing data to the centromere assemblies from diverse primate species shows uniform read depth, indicating a lack of large structural errors. The human genome is HG00733.

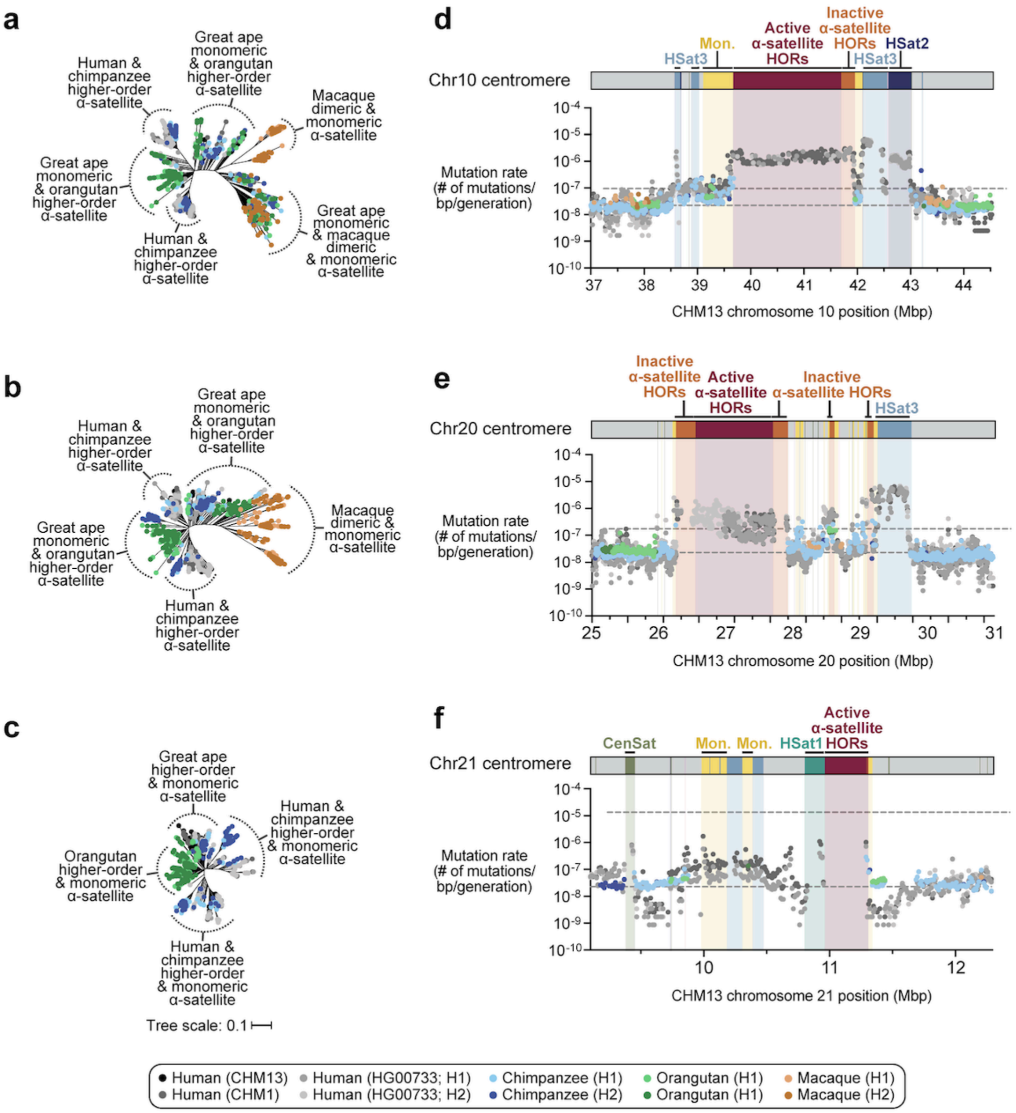


**Extended Data Figure 17. Centromeres evolve with different evolutionary trajectories and mutation rates. a-c)** Phylogenetic trees of α-satellite monomers derived from the human, chimpanzee, orangutan, and macaque chromosome **a)** 10, **b)** 20, and **c)** 21 centromeric regions. **d-f)** Plot showing the mutation rate of the chromosome **d)** 10, **e)** 20, and **f)** 21 centromeric regions. Individual data points from 10-kbp pairwise sequence alignments are shown.

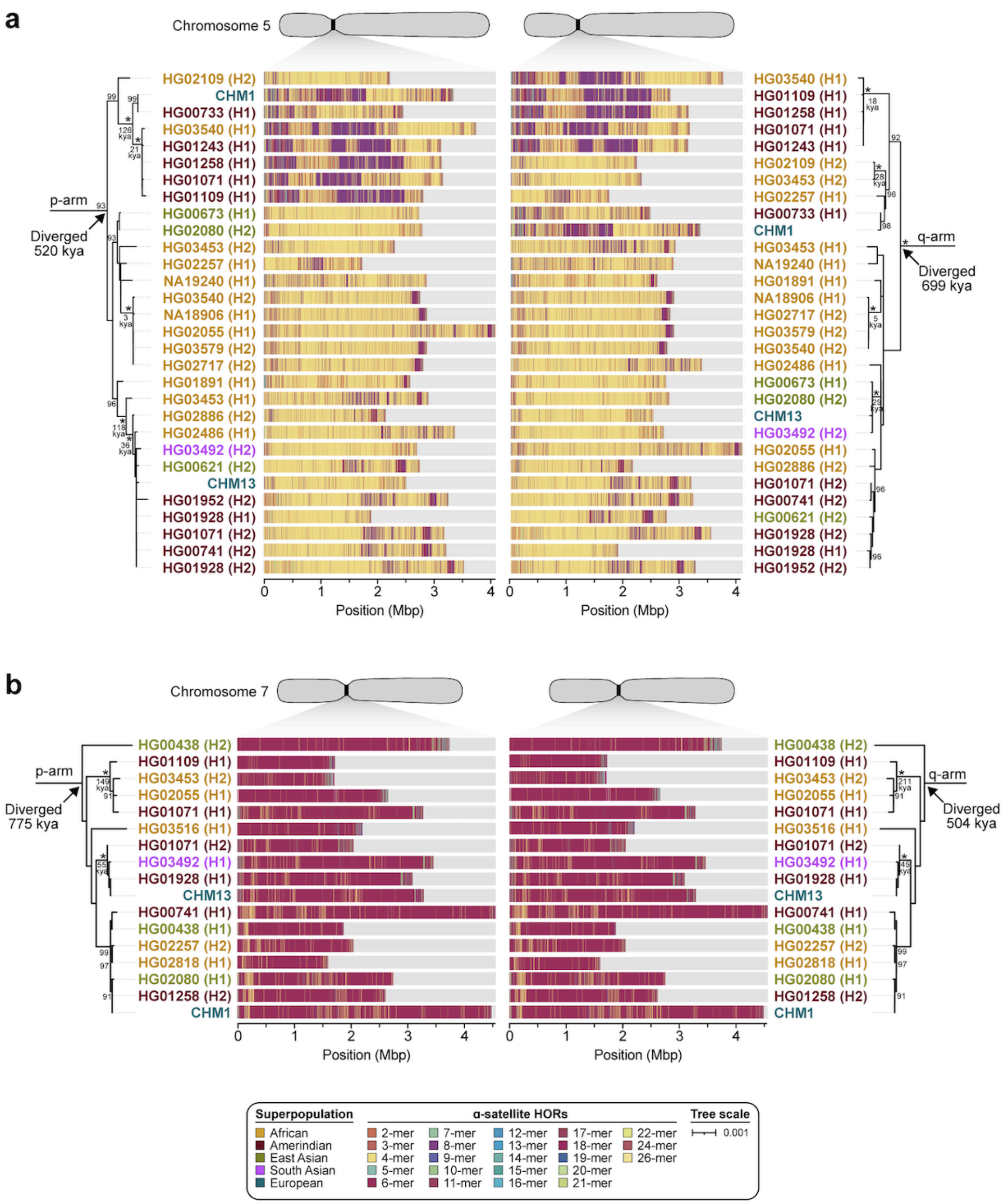


**Extended Data Figure 18. Phylogenetic reconstruction of human chromosome 5 and 7 centromeric haplotypes. a,b)** Phylogenetic trees showing the evolutionary relationship and estimated divergence times of completely and accurately assembled **a)** *D5Z2* α-satellite HOR arrays and **b)** *D7Z1* α-satellite HOR arrays from CHM1, CHM13, and diverse human samples (generated by the HPRC[9] and HGSVC[10]). The trees were generated from 20-kbp segments in the monomeric α-satellite or unique sequence regions on the p- (left) and q- (right) arms.

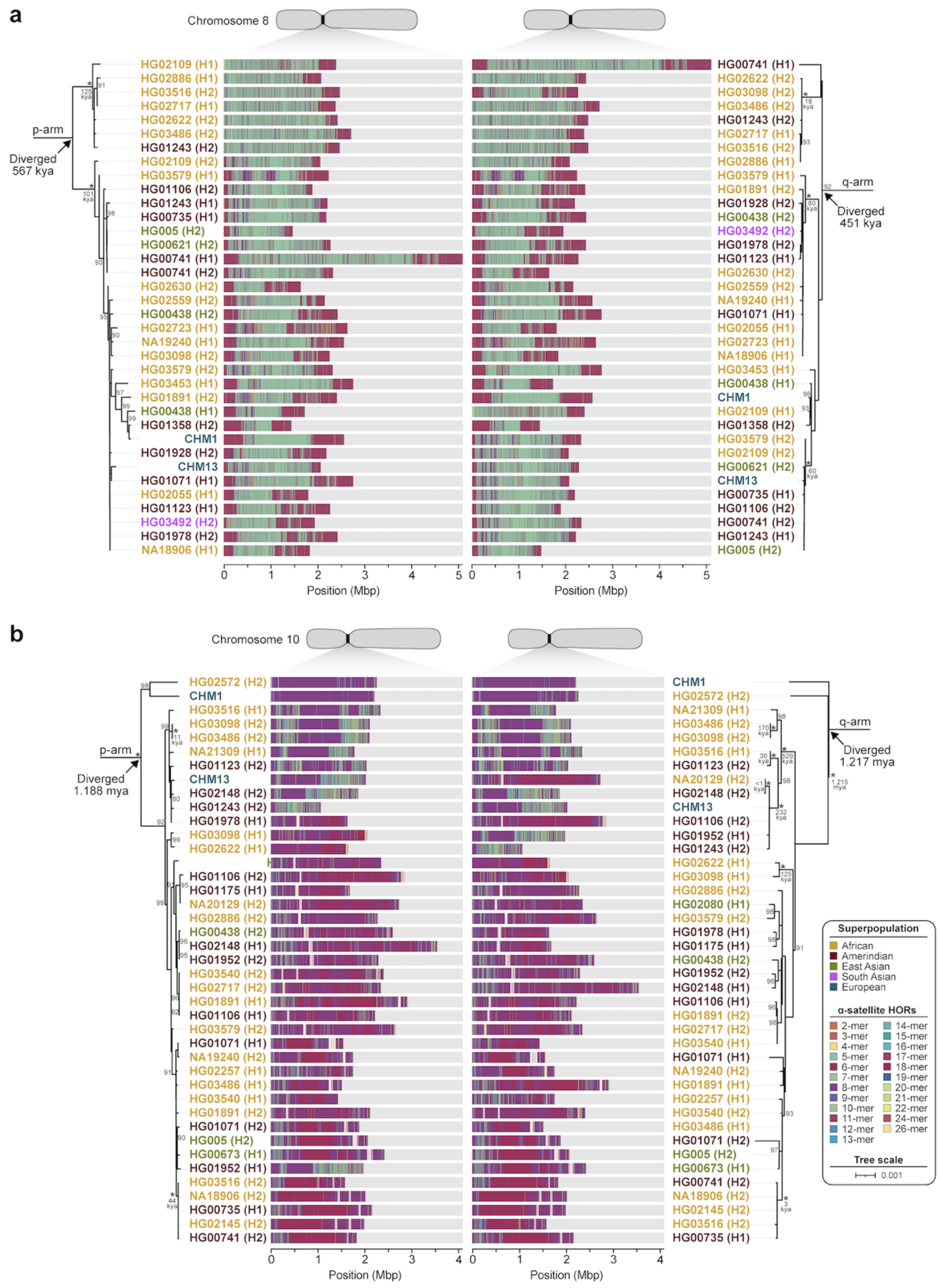


**Extended Data Figure 19. Phylogenetic reconstruction of human chromosome 8 and 10 centromeric haplotypes. a,b)** Phylogenetic trees showing the evolutionary relationship and estimated divergence times of completely and accurately assembled **a)** *D8Z2* α-satellite HOR arrays and **b)** *D10Z1* α-satellite HOR arrays from CHM1, CHM13, and diverse human samples (generated by the HPRC[9] and HGSVC[10]). The trees were generated from 20-kbp segments in the monomeric α-satellite or unique sequence regions on the p- (left) and q- (right) arms.

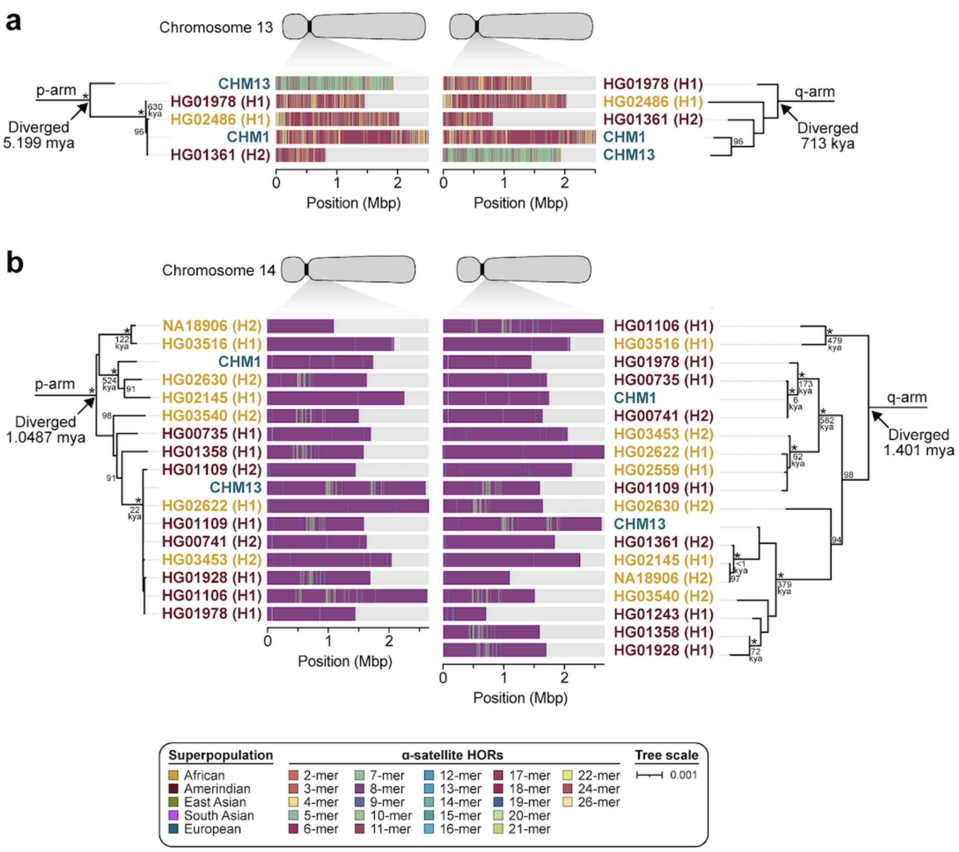


**Extended Data Figure 20. Phylogenetic reconstruction of human chromosome 13 and 14 centromeric haplotypes. a,b)** Phylogenetic trees showing the evolutionary relationship and estimated divergence times of completely and accurately assembled **a)** *D13Z2* α-satellite HOR arrays and **b)** *D14Z9* α-satellite HOR arrays from CHM1, CHM13, and diverse human samples (generated by the HPRC[9] and HGSVC[10]). The trees were generated from 20-kbp segments in the monomeric α-satellite or unique sequence regions on the p- (left) and q- (right) arms.

### EXTENDED DATA TABLES

**Extended Data Table 1. Statistics of long-read sequencing datasets and genome assemblies.** See accompanying Excel file.

**Extended Data Table 2. Support for CHM1 and CHM13 centromere assemblies from 56 diverse human genomes sequenced by the HPRC and HGSVC.** See accompanying Excel file.

**Extended Data Table 3. Three alignment strategies for assessing centromere sequence identity.** See accompanying Excel file.

**Extended Data Table 4. Sequence identity calculated from full contig alignments between CHM1 and CHM13 centromeres.** See accompanying Excel file.

**Extended Data Table 5. Sequence identity calculated from alignments of 10-kbp segments between the CHM1 and CHM13 centromeres.** See accompanying Excel file.

**Extended Data Table 6. Sequence identity and alignment statistics of centromeric α-satellite HOR arrays from CHM1, CHM13, and 56 diverse human genomes.** See accompanying Excel file.

**Extended Data Table 7. Quantification of the genetic and epigenetic variation of all CHM1 and CHM13 centromeres.** See accompanying Excel file.

**Extended Data Table 8. Catalog of all α-satellite HOR variants in the CHM1 and CHM13 centromeres.** See accompanying Excel file.

**Extended Data Table 9. Quantification of changes in bases, α-satellite monomers, and α-satellite HORs among centromeric arrays with the same monophyletic origin.** See accompanying Excel file.

**Extended Data Table 10. Datasets generated and/or used in this study.** See accompanying Excel file.

### EXTENDED DATA REFERENCES
