## Supplementary Information for "The variation and evolution of complete human centromeres"

**This PDF file includes:**

1. **Supplementary Notes 1 and 2**
2. **References**

### SUPPLEMENTARY NOTES

**Supplementary Note 1. Analysis of the CHM1 karyotype.** We first assessed the karyotype of the CHM1 genome using three orthogonal methods: Giemsa staining, spectral karyotyping, and single-cell sequencing of template DNA strands (Strand-seq; **Extended Data Figs. 2,3**). All three methods indicate that the CHM1 cell population is biclonal, with approximately 71% of cells existing in a diploid or near-diploid state and approximately 29% of cells existing in a tetraploid or near-tetraploid state (**Extended Data Figs. 2a,b**). Both diploid/near-diploid and tetraploid/near-tetraploid cells have multiple chromosomal rearrangements, including a translocation between chromosomes 4q35.1 and 11q24.3 (**Extended Data Figs. 2c,d**), a translocation between chromosomes 16q23.3 to 17q25.3 (**Extended Data Figs. 2e,f**), and a loss of chromosome 17 or the chromosome 17 p-arm in a subset of cells (**Extended Data Fig. 2f**). The translocation between chromosomes 4q35.1/11q24.3 is accompanied by a complete deletion of *STOX2* and *ADAMTS15* and partial deletion of the *ADAMTS8* (**Extended Data Fig. 3c**). *ADAMTS15* is predicted to act as a tumor suppressor gene in breast and colorectal cancer[1,2]. Additionally, the translocation between chromosomes 16q23.3 to 17q25.3 results in a novel gene fusion between *CDH13* and *RPTOR* (**Extended Data Fig. 3d**), which are both associated with cancer[3–6] and may contribute to the observed karyotype of the CHM1 cell line.

**Supplementary Note 2. Loss of two distinct chromosomal regions in the CHM1 cell line.** Mapping of native PacBio HiFi and ONT long-read sequencing data to the CHM1 centromere assemblies reveals a reduction in coverage on the p-arm-proximal side of the chromosome 17 centromere (**Extended Data Fig. 4**), consistent with the loss of the p-arm in a subset of cells (**Extended Data Fig. 2f**). It also reveals a reduction in coverage over a 631-kbp region in the *D13Z2* α-satellite higher-order repeat (HOR) array on chromosome 13 (**Extended Data Fig. 4**), indicating this region is deleted in a subset of cells.

### SUPPLEMENTARY INFORMATION REFERENCES
